# Slx4 and Fun30/SMARCAD1 coordinate S-phase checkpoint regulation and replication fork protection in response to Top1-DNA crosslinks

**DOI:** 10.1101/2025.07.28.667195

**Authors:** Mathilde Courtes, Thierry Boissière, Antoine Barthe, Philippe Pasero, Benjamin Pardo

## Abstract

Replication stress is a major driver of genomic instability and is implicated in the development of diseases such as cancer. It triggers the S-phase checkpoint, a signaling pathway that coordinates the handling of replication obstacles with cell cycle progression. One prominent source of replication stress is the formation of DNA-protein crosslinks on the template, such as those induced by DNA topoisomerase I poisoning by camptothecin (CPT). In this study, we investigated how the S-phase checkpoint responds to CPT-induced replication stress. We show that both activation and timely deactivation of checkpoint signaling are critical for DNA replication completion and cell viability. Using a locus-specific approach, we found that checkpoint signaling is actively dampened at lesion sites. Mechanistically, this attenuation involves the displacement of the checkpoint mediator Rad9 by the DNA repair factors Slx4 and Fun30. This local dampening not only promotes cell cycle progression, but also permits Exo1-dependent resection of replication forks stalled by Top1-DNA crosslinks. Controlled resection, in turn, allows homologous recombination factors to access and stabilize the forks, preventing their degradation. In conclusion, we propose that local checkpoint dampening by Slx4 and Fun30 at replication stress sites is a critical mechanism that promotes replication completion and preserves genome stability.

**Graphical abstract:** 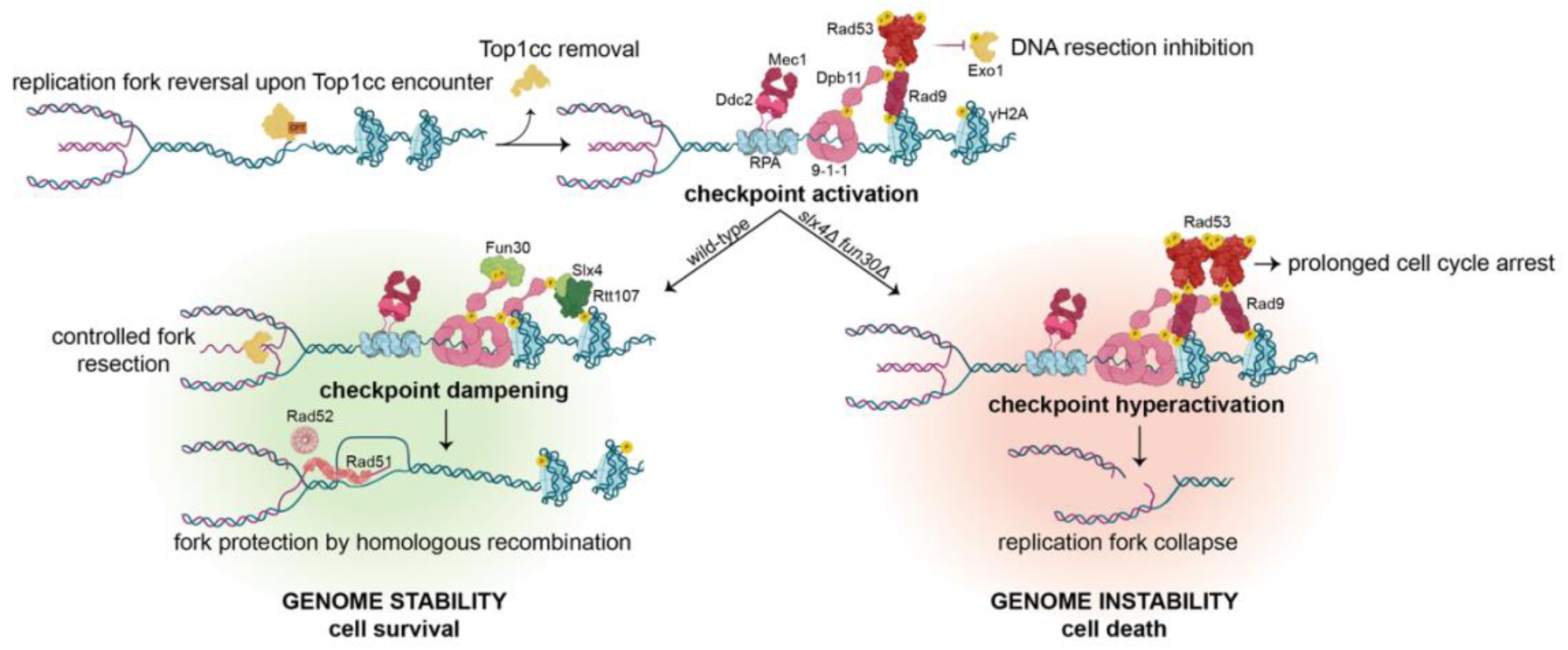

## Introduction

During the S phase of the cell cycle, replication forks can encounter obstacles on the DNA template that impede their progression, a condition commonly referred to as replication stress. Prolonged fork arrest can lead to their collapse, resulting in DNA lesions that may cause mutations or chromosomal rearrangements if misrepaired. Therefore, replication stress is considered a major driver of genomic instability. One physiological event that generates replication stress is the persistence of Top1 cleavage complexes (Top1ccs) on the DNA template. DNA topoisomerase 1 (Top1) is an enzyme that catalyzes the unwinding of supercoiled DNA through a concerted mechanism of transient DNA strand cleavage and religation. Top1ccs are transient intermediates in the Top1 catalytic cycle, formed when Top1 covalently binds to the 3’ end of a single-stranded DNA break via a tyrosine residue in its active site. Top1ccs can become trapped at naturally occurring DNA lesions, such as nicks, gaps, damaged bases or abasic sites (1, 2).

Top1ccs trapping occurs with such high frequency that specific repair enzymes have evolved to remove it. The proteasome or SPRTN protease is targeted to degrade abortive Top1ccs (3), and tyrosyl-DNA phosphodiesterase 1 (TDP1) can specifically hydrolyze the phosphotyrosine bond that binds Top1 to DNA (4). The combined loss of SPRTN and TDP1 results in near-synthetic lethality, indicating that these factors have redundant roles in Top1cc removal (5, 6). Despite the existence of these pathways, highly replicating cells are particularly affected by Top1ccs (7).

The persistence of Top1ccs is also enhanced by camptothecin (CPT), a drug that binds the phosphotyrosine bond between Top1 and DNA, blocking religation and subsequent release of the topoisomerase (8). CPT derivatives such as topotecan and irinotecan are commonly used in chemotherapy to induce replication stress, selectively targeting highly proliferative cells. Replication stress induced by CPT is thought to arise from the conversion of single-stranded DNA breaks at Top1ccs into single-ended double-strand breaks (DSBs) when replicative helicases separate the parental DNA strands (9). However, neither we nor others have observed replication-dependent DSBs in CPT-treated wild-type yeast cells. Instead, we favor a model in which the replication impediment results from the reversal of replication forks encountering Top1ccs (10–12). Fork reversal involves the pairing of newly synthesized DNA strands and the rewinding of parental DNA, causing the fork to move backwards, which would provide more space for Top1cc removal. We recently proposed that homologous recombination (HR) factors, typically involved in DSB repair, are recruited to reversed forks to protect them from degradation until they merge with converging replication forks (11). Without such protection, reversed forks become vulnerable to nucleolytic cleavage, leading to genomic instability (11, 13, 14).

Cellular responses to replication stress are tightly coordinated by the S-phase checkpoint. This pathway safeguards DNA replication by preventing the activation of late replication origins and slowing the elongation of active forks, while preserving their stability and functionality (15–20). In addition, it arrests the cell cycle at the G2/M transition to ensure complete DNA synthesis and repair before entry into mitosis (21, 22). The S-phase checkpoint is activated by the accumulation of single-stranded DNA (ssDNA) at sites of DNA damage or stressed forks. This results either from nuclease activity or from uncoupling between the replicative helicase and DNA polymerases, or between leading and lagging strand synthesis. ssDNA stretches recruit the sensor kinase ATR through its partner ATRIP. ATR activation is further enhanced by TopBP1, which binds single-stranded/double-stranded 5’ DNA junctions via its interaction with the 9-1-1 complex (23, 24). Once activated, ATR initiates a signaling cascade that transmits the checkpoint signal to the effector kinase CHK1 via mediator proteins such as CLASPIN, 53BP1 and MDC1 (23, 25, 26).

Conserved signaling pathways have also been described in the budding yeast *Saccharomyces cerevisiae*. Extensive studies have distinguished two distinct branches within the S-phase checkpoint: the DNA replication checkpoint (DRC) and DNA damage checkpoint (DDC) (27). These pathways work together to generate an integrated cellular response to replication stress but differ in their activation trigger, spatial regulation and activation kinetics, primarily determined by the mediator used by Mec1^ATR^ to activate Rad53^CHK1/CHK2^ (18, 28). DRC activation is mediated by Mrc1, the yeast homolog of CLASPIN, which is a replisome component sensitive to ssDNA accumulation at replication forks. In contrast, DDC activation depends on Rad9, the functional ortholog of 53BP1 and MDC1. Rad9^53BP1/MDC1^ is recruited to chromatin via three distinct platforms: (*i*) Rad9 binds methylated histone H3 on K79. Methylation of H3K79 by Dot1 is not induced by DNA damage, but is associated with cell cycle progression. (*ii*) Upon DNA damage, Rad9 can bind to γH2A(X), the phosphorylated form of H2A histone on S129 by Mec1^ATR^ or Tel1^ATM^. Mec1 also phosphorylates the 9-1-1 complex, allowing further recruitment of the multi-BRCT domain protein Dpb11^TOPBP1^. (*iii*) Phosphorylation of Rad9 by the cyclin-dependent kinase (CDK) Cdc28 allows its binding to Dpb11, making this last interaction cell cycle- and damage-dependent. Rad9 can simultaneously interact with γH2A(X) and Dpb11, reinforcing its localization to DNA damage sites, even though each independent interaction is sufficient for Rad53 recruitment and activation. The full activation of Rad53 is primed by Mec1 phosphorylation and is followed with extensive trans-autophosphorylation. This promotes the dissociation of Rad53 from chromatin and dissemination of the checkpoint signal throughout the nucleus.

Just as checkpoint activation is essential for surviving replication stress, timely checkpoint deactivation is equally important for cell cycle resumption and cell growth (29). Rather than being driven by Rad53 degradation, checkpoint deactivation involves the restauration of Rad53 in its unphosphorylated form, either through direct dephosphorylation or by downregulating its phosphorylation. Although multiple phosphatases act on the pool of activated Rad53, their activity is not sufficient to ensure a timely downregulation of the checkpoint response. Pioneering work by Ohouo et al (2013) identified a phosphatase-independent mechanism of checkpoint deactivation, termed checkpoint dampening, which acts locally at sites of damage. They showed that the DNA repair scaffold formed by Rtt107 and Slx4 can bind γH2A(X) through Rtt107 and Dpb11 through Slx4 in a manner that is mutually exclusive with Rad9 binding. This competitive binding leads to Rad9 displacement from chromatin, thereby reducing Rad53 activation. This dampening mechanism has been studied in response to DSBs and methyl methanesulfonate (MMS)-induced replication stress (30–32). Whether this dampening mechanism also operates at CPT-induced reversed forks and what are the consequences of defective dampening at the molecular level remains to be investigated.

In this study, we characterized the checkpoint response to CPT-induced replication stress and showed that both activation and subsequent deactivation of the checkpoint are critical for cell viability. We found that Rad53 activation is markedly attenuated in response to CPT. This attenuation involves the displacement of Rad9 from DNA damage sites by the Slx4-Rtt107 complex, as well as by the chromatin remodeler Fun30, independently of their canonical roles in DNA repair. Loss of this checkpoint dampening mechanisms led to checkpoint hyperactivation and inhibition of cell cycle progression. Importantly, it also impaired the Exo1-dependent resection of nascent DNA at stalled replication forks, a critical step for the initiation of homologous recombination. We conclude that checkpoint suppression by Slx4-Rtt107 and Fun30 is a key regulatory mechanism that promotes the HR-mediated protection of replication forks encountering Top1-DNA crosslinks.

## Materials and Methods

### Biological Ressources

All *Saccharomyces cerevisiae* strains used are haploid and are derived from W303 background (*his3-11, 15 leu2-3, 112, trp1-1, ura3-1, ade2-1, can1-100*) and corrected for *rad5-535* mutation, unless specifically stated. All strains were obtained either by PCR transformation (33) or by genetic crosses (verified by individual tetrad analysis). Modifications were verified by PCR, drug sensitivity analysis and western blot when needed. Detailed strain genotypes can be found in **Table 1**.

**Table 1.** Yeast strains used in this study.

| **Table 1. Yeast strains used in this study** |  |
| --- | --- |
| Strain N° | Genotype |
| PP4216/5319 | *MATa bar1Δ* |
| PP4253/4254 | *MATa bar1Δ Mms4-His6-Flag10-KAN* |
| PP4271 | *MATa bar1Δ URA3::GPD-TK7 Mms4-His6-Flag10-KAN* |
| PP4241 | *MATa, bar1Δ URA3::GPD-TK7* |
| **Checkpoint** |  |
| PP4271/4272 | *MATa bar1Δ URA3::GPD-TK7 Mms4-His6-Flag10-KAN* |
| PP4369/4370 | *MATa, bar1Δ::LEU2 mec1Δ::TRP1, sml1Δ::HIS3 Mms4-His6-Flag10-KAN* |
| PP4301/4302 | *MATa bar1Δ URA3::GPD-TK7 Mms4-His6-Flag10-KAN rad9ΔklTRP1* |
| PP4303/4304 | *MATa bar1Δ URA3::GPD-TK7 Mms4-His6-Flag10-KAN mrc1ΔklTRP1* |
| PP3962/3963 | *MATa bar1Δ mrc1Δ::HIS5, mrc1-AQ::LEU2* |
| PP5174/5175 | *MATa, URA3::GPD-TK, mrc1Δ::HIS, rad9Δ::LEU2, sml1Δ::KAN* |
| PP944/945 | *Mat a, URA3::GPD-TK, sml1::HIS rad53::KAN* |
| PP3996/3997 | *MATa bar1Δ URA3::GPD-TK7 rad53-11* |
| PP5128/5129 | *MATa, bar1Δ::NAT, URA3::GPD-TK7 chk1Δ::LEU2* |
| PP5134/5135 | *MATa, bar1Δ::NAT, URA3::GPD-TK7 mec2-1 (rad53-11), chk1Δ::LEU2* |
| **Dampening** |  |
| PP4179/4180 | *MATa bar1Δ slx4Δ::klLEU2 Mms4-His6-Flag10-KAN* |
| PP4181/4182 | *MATa bar1Δ slx4Δ::klLEU2 fun30Δ::NAT Mms4-His6-Flag10-KAN* |
| PP4469/4470 | *MATa, bar1Δ URA3::GPD-TK7 slx4Δ::klLEU2* |
| PP4653/4654 | *MATa, bar1Δ URA3::GPD-TK7 fun30Δ::KanMX4* |
| PP4655/4656 | *MATa, bar1Δ URA3::GPD-TK7 slx4Δ::klLEU2 fun30Δ::KanMX4* |
| PP4030/4031 | *MATa, bar1Δ URA3::GPD-TK7 slx4Δ::klLEU2 rad9Δ::klTRP1* |
| PP4084/4085 | *MATalpha/a fun30Δ::KAN rad9Δ::TRP1* |
| PP4026/4027 | *MATa bar1Δ URA3::GPD-TK7 fun30Δ::KanMX4 fun30-SSAA::TRP1* |
| PP4028/4029 | *MATa bar1Δ URA3::GPD-TK7 slx4Δ::klLEU2 fun30Δ::KanMX4 fun30-SSAA::TRP1* |
| PP7097/7098 | *MATa bar1Δ fun30Δ::KAN fun30-K603R::TRP1* |
| PP7222/7223 | *MATa bar1Δ fun30Δ::KAN fun30-K603R-SSAA::TRP1* |
| PP5344/5345 | *MATa bar1Δ rad9ΔHIS rad9-STAA::TRP1* |
| PP5348/5349 | *MATa bar1Δ rad9ΔHIS rad9-STAA::TRP1 slx4Δ::klLEU2* |
| PP5352/53 | *MATa bar1Δ rad9ΔHIS rad9-STAA::TRP1 fun30Δ::KAN* |
| PP5356/5357 | *MATa bar1Δ rad9ΔHIS rad9-STAA::TRP1 slx4Δ::klLEU2 fun30Δ::KAN* |
| PP6153/6154 | *MATa bar1Δ rad9ΔHIS rad9-STAA::TRP1 fun30Δ::KAN fun30-SSAA::URA3* |
| PP6155/6156 | *MATa bar1Δ rad9ΔHIS rad9-STAA::TRP1 slx4Δ::klLEU2 fun30Δ::KAN fun30-SSAA::URA3* |
| Recombination |  |
| PP4357/4358 | *MATa, bar1Δ::NAT URA3::GPD-TK7 mus81ΔKAN URA3::GPD-TK(5x), AUR1c::ADH-hENT1* |
| PP4347/4348 | *MATa bar1Δ Mms4-His6-Flag10-KAN mus81ΔklURA3* |
| PP4183/4184 | *MATa bar1Δ mus81Δ::klURA3 slx4Δ::klLEU2* |
| PP4185/4186 | *MATa bar1Δ mus81Δ::klURA3 fun30Δ::NAT* |
| PP4187/4188 | *MATa bar1Δ mus81Δ::klURA3 slx4Δ::klLEU2 fun30Δ::NAT* |
| PP4243/4244 | *MATa, bar1Δ::NAT URA3::GPD-TK7 mus81ΔKAN* |
| PP4521 | *MATa, bar1Δ URA3::GPD-TK7 slx4Δ::klLEU2 mus81Δ::NAT* |
| PP7107/7108 | *MATa bar1Δ mus81Δ::klURA3 rad1Δ::HYG slx1ΔHIS* |
| PP5873/(5874) | *MATa, (rad5-535), Rad52-YFP Rfa1-CFP mCHERRY1-Pus1::URA3 YEN1-ON-Myc9::KAN* |
| PP5727/5728 | *MATa, Rad52-YFP Rfa1-CFP mCHERRY1-Pus1::URA3 YEN1-ON-Myc9::KAN slx4Δ::NAT* |
| PP5978/5979 | *MATa, Rad52-YFP Rfa1-CFP mCHERRY1-Pus1::URA3 YEN1-ON-Myc9::KAN slx4Δ::NAT fun30Δ::HYG* |
| PP4407 | *MATa bar1Δ rad52Δ::klLEU2* |
| Fluorescence |  |
| PP3558/3559 | *MATa, rad5-535, Rad52-YFP Rfa1-CFP mCHERRY1-Pus1::URA3* |
| PP3564/3565 | *MATa, rad5-535, Rad52-YFP Rfa1-CFP mCHERRY1-Pus1::URA3 slx4::NAT* |
| PP4824/4825 | *MATa, rad5-535, RAD52-YFP RFA1-CFP mCHERRY-PUS1::URA3 fun30Δ::KanMX4* |
| PP4835/4836 | *MATa, rad5-535, RAD52-YFP RFA1-CFP mCHERRY-PUS1::URA3 slx4Δ::NAT fun30Δ::KanMX4* |
| PP4826/4827 | *MATa, rad5-535, RAD52-YFP RFA1-CFP mCHERRY-PUS1::URA3 fun30Δ::KanMX4 fun30-SSAA::TRP1* |
| PP4833/4834 | *MATa, rad5-535, RAD52-YFP RFA1-CFP mCHERRY-PUS1::URA3 slx4Δ::NAT fun30Δ::KanMX4 fun30-SSAA::TRP1* |
| Resection |  |
| PP6393/6394 | *MATa bar1Δ pRS424-empty-TRP1* |
| PP6395/6396 | *MATa bar1Δ slx4Δ::klLEU2 pRS424-empty-TRP1* |
| PP6397/6398 | *MATa bar1Δ fun30Δ::KAN pRS424-empty-TRP1* |
| PP6399/6400 | *MATa bar1Δ slx4Δ::klLEU2 fun30Δ::KAN pRS424-empty-TRP1* |
| PP6401/6402 | *MATa bar1Δ pRS424-EXO1-TRP1* |
| PP6403/6404 | *MATa bar1Δ slx4Δ::klLEU2 pRS424-EXO1-TRP1* |
| PP6405/6406 | *MATa bar1Δ fun30Δ::KAN pRS424-EXO1-TRP1* |
| PP6407/6408 | *MATa bar1Δ slx4Δ::klLEU2 fun30Δ::KAN pRS424-EXO1-TRP1* |
| PP6409/6410 | *MATa bar1Δ pRS424-exo1-D173A-TRP1* |
| PP6411/6412 | *MATa bar1Δ slx4Δ::klLEU2 pRS424-exo1-D173A-TRP1* |
| PP6413/6414 | *MATa bar1Δ fun30Δ::KAN pRS424-exo1-D173A-TRP1* |
| PP6415/6416 | *MATa bar1Δ slx4Δ::klLEU2 fun30Δ::KAN pRS424-exo1-D173A-TRP1* |
| PP6582/6583 | *MATa bar1Δ rad9ΔHIS rad9-STAA:URA3 prs424-empty-TRP1* |
| PP6588/6589 | *MATa bar1Δ rad9ΔHIS rad9-STAA:URA3 slx4Δ::klLEU2 fun30Δ::NAT prs424-empty-TRP1* |
| PP6590/6591 | *MATa bar1Δ rad9ΔHIS rad9-STAA:URA3 prs424-EXO1-TRP1* |
| PP6596/6597 | *MATa bar1Δ rad9ΔHIS rad9-STAA:URA3 slx4Δ::klLEU2 fun30Δ::NAT prs424-EXO1-TRP1* |
| PP6972 | *WT MATa bar1Δ EXO1-Myc13-KAN* |
| PP6978 | *WT MATa bar1Δ exo1-23A-Myc13-KAN* |
| PP6976/6977 | *WT MATa bar1Δ EXO1-Myc13-KAN slx4ΔklLEU2 fun30ΔNAT* |
| PP6982/6983 | *WT MATa bar1Δ exo1-23A-Myc13-KAN slx4ΔklLEU2 fun30ΔNAT* |
| PP5199 | *MATa, bar1Δ exo1::NAT, sgs1::LEU2 ARS608::HIS3, ARS609::TRP1, ARS610::HYG mCHERRY1-Pus1::URA3* |
| eRFB system |  |
| PP4678/4679 | *MATa, bar1::LEU2 Top1-Flag3::TRP1, tdp1::NAT* |
| PP4680/4681 | *MATalpha, ura3-1::ADH1-OsTIR1-2-9Myc::URA3 wss1-miniAID-9myc::HYG* |
| PP4691/4692 | *MATalpha/a, bar1::LEU2 ura3-1::ADH1-OsTIR1-2-9Myc::URA3 wss1-miniAID-9myc::HYG tdp1::NAT Top1-Flag3::TRP1* |
| PP3497/3498 | *MATa, bar1Δ ura3-1::ADH1-OsTIR1-2-9Myc::URA3 wss1-miniAID-9myc::HYG tdp1::NAT pdr1::spHIS5 MRP1-KAN-RFB-Inverted-PAL1 Top1-Flag3::TRP1* |
| PP6298/6299 | *MATa, bar1Δ ade2-1::OsTIR1-9myc::ADE2 wss1-miniAID-9myc::HYG tdp1::NAT pdr1::spHIS5 MRP1-KAN-RFB-Inverted-PAL1 Top1-Flag3::TRP1* |
| PP6474/6475 | *MATa, bar1Δ ade2-1::OsTIR1-9myc::ADE2 wss1-miniAID-9myc::HYG tdp1::NAT pdr1::spHIS5 MRP1-KAN-PAL1 Top1-Flag3::TRP1* |
| PP6643/6644 | *MATa, bar1Δ ade2-1::OsTIR1-9myc::ADE2 wss1-miniAID-9myc::HYG tdp1::NAT pdr1::spHIS5 MRP1-KAN-RFB-Inverted-PAL1 Top1-Flag3::TRP1 fun30::klURA3* |
| PP6645/6646 | *MATa, bar1Δ ade2-1::OsTIR1-9myc::ADE2 wss1-miniAID-9myc::HYG tdp1::NAT pdr1::spHIS5 MRP1-KAN-RFB-Inverted-PAL1 Top1-Flag3::TRP1 slx4::klURA3* |
| PP6691/92 | *MATa, bar1Δ ade2-1::OsTIR1-9myc::ADE2 wss1-miniAID-9myc::HYG tdp1::NAT pdr1::spHIS5 MRP1-KAN-RFB-Inverted-PAL1 Top1-Flag3::TRP1 slx4Δ::klLEU2* |
| PP6693/6694 | *MATa, bar1Δ ade2-1::OsTIR1-9myc:ADE2 wss1-miniAID-9myc::HYG tdp1::NAT pdr1::spHIS5 MRP1-KAN-RFB-Inverted-PAL1 Top1-Flag3::TRP1 fun30Δ::klURA3 slx4Δ::klLEU2* |

Time courses were initiated from YPAD overnight mid-log cultures and normalized to 7.10^6^ cells/ml after counting using a CASY^®^ (OLS system) flow cytometer. α-factor was added to the culture medium to block cells in G1. Unless stated otherwise, G1-blocked cells were washed with MPD +SDS medium (0.17% yeast nitrogen base, 0.1% L-proline, 2% glucose, and 0.003% SDS), resuspended in MPD +SDS medium supplemented with α-factor and DMSO or CPT, incubated for 1 more hour, and released into S phase by addition of pronase (50 µg/ml) to the culture medium.

### Statistical Analyses

All statistical tests and numbers of biological replicates are listed in the figure legends. Mean±SEM are displayed for the concerned experiments, and sample size was not predetermined using any statistical method. All statistical tests were performed with GraphPad Prism 10.

### Protein extraction, electrophoresis and immunoblotting

We have analyzed the presence and/or the phosphorylation state of our proteins of interest by detecting an electrophoretic mobility shift by western blot.

5mL of the desired cultures (∼1.10^7^ cells/ml after synchronization) were collected in tubes containing 50µL of 10% Sodium Azide (for 0,1% final) and kept on ice until processing. Proteins were extracted from cell pellets with TCA 20% and Zirconium glass beads after 15min vortexing at 2400 rpm on a VXR basic Vibrax^®^ apparatus. Extracts were resolved by SDS-PAGE in precast 3-8% polyacrylamide gels (Invitrogen), transferred with Trans-Blot^®^ Turbo^TM^ system (Bio-Rad) to nitrocellulose membranes and probed with α-Rad53 (gift from Corrado Santocanale), α-MYC (sc-40, SantaCruz Technology) or α-FLAG (F3165, Sigma) antibodies in TBS 1x +Tween 0,1% +BSA 1% +NaN3 0,02%. Secondary rabbit and mouse antibodies coupled to horseradish peroxidase were used for revelation with the SuperSignal West Pico Plus chemiluminescent substrate and detected with a ChemiDoc apparatus (Bio-rad). Two to four independent biological replicates were performed.

To quantify Rad53 protein phosphorylation, the box tool from Image J software was overlaid onto Rad53 total signal. Area under the curve of the derived plots was retrieved. Separation between highly-phosphorylated and low-phosphorylated Rad53 was established as the lowest point between the two resulting pics. A line was drawn at this location and the area under the curve of the phosphorylated form was retrieved. Raw area values were analyzed in GraphPad Prism for quantification and statistical analyses. Plots show percentage of phosphorylated Rad53 over total amount. At least three independent biological replicates have been performed for each assay.

When specified, Phos-tag™ reagent (Wako chemicals, AAL-107) and fresh MnCl2 were added to 7.5% sodium dodecyl sulphate-polyacrylamide gel electrophoresis (SDS-PAGE) gels to a final concentration of 5 µM and 100 µM, respectively. Extracts were resolved at constant voltage 150V per gel in a Mini-protean III (Bio-Rad) for 150 min. After the electrophoresis, the gel was incubated in standard methanol-based transfer buffer with ethylenediaminetetraacetic acid (EDTA) at 0.1M for 30 min. Then, the gel was transferred to a PVDF membrane (Immobilon-P, Millipore; previously activated with a quick rinse of methanol 100% followed by washes with water and transfer buffer) in a Mini-Transfer system (Bio-Rad) at 30V overnight with ice block and stirring. Exo1-MYC was detected using the mouse monoclonal α-MYC antibody (sc-40, SantaCruz Technology).

### Cell cycle progression analyses

Flow cytometry was used to follow the cell-cycle progression, by measuring the DNA content going from one content (1C) to two (2C) as genome duplication occurs.

Like previously mentioned, YPAD overnight mid-log cultures were normalized to 7.10^6^ cells/ml and blocked in G1 with αfactor addition for two hours at 30°C. Cells were then washed of the YPAD medium and resuspended in a permeabilizing medium (11) containing either DMSO or 50µM CPT still in the presence of αfactor for one more hour of incubation. Cells were synchronously released into S-phase via pronase (50µG/mL) addition to the culture medium.

Samples were harvested and fixed with 100% ethanol for subsequent flow cytometry analysis. Fixed samples were centrifuged for 1min at 15,000 rpm, resuspended in 50mM sodium citrate buffer and sequentially treated with RNAse A (0,2mg/mL, Qiagen 76254; 2h at 50°C or O/N at 37°C) and Proteinase K (0,24mg/mL, Sigma, P6556; 1h at 50°C). Aggregates of cells were dissociated by sonication and stained with propidium iodide (PI, 4µg/mL, Sigma) for at least 2 hours in darkness. PI fluorescence, reflecting DNA content, was measured by flow cytometry. Gate was fixed to 10.000 events. Data was acquired with the MACSQuant analyzer (Miltenyi Biotech) and analyzed with FlowJo software. Two to four independent biological replicates were performed for each analysis.

Dotted lines were drawn from the 1C and 2C population peaks in the corresponding wild-type condition and reported as such for other strains. In this way, we determined the extent of S phase, which ended when the bulk cell population reached the 2C dotted line, and the extent of G2/M phase, which ended when the 2C population dropped. For ease of reading, each cycle step determined by this method (asynchronous, G1 block, S and G2/M) is highlighted with a specific color when needed (pink, green, blue and orange, respectively).

### Camptothecin (CPT) sensitivity assays (Drop Tests)

Cells were counted and concentrated to 2.10^8^ cells/mL. 4µL of 10-fold serial dilutions were spotted on rich YPAD or SC plates +/- CPT ((S)-(+)-Camptothecin, Sigma #C9911). Plates were incubated respectively 2 or 3 days at 30°C.

### Pulsed-field gel electrophoresis (PFGE) and Southern Blot

We have used PFGE to separate individual yeast chromosomes according to their size thanks to periodical changes in the direction of the electric field. DNA was purified in agarose plugs as described (34) and loaded on agarose gels. Effective chromosome separation was achieved by changing electric pulses direction (interval from 100 to 10 s (logarithmic), angle from 120 to 110° (linear), and voltage from 200 to 150 V (logarithmic)) during 24h in TBE 0.3× at 13°C using a Rotaphor^®^ apparatus (Biometra). The gel was stained with ethidium bromide before being transferred to a nitrocellulose membrane Hybond XL (GE Healthcare) for Southern blotting. The membrane was hybridized using a P^32^ labelled DNA radioactive probe specific for the *ACT1* gene on chromosome VI. Two to three independent biological replicates were performed.

### Foci formation microscopy analysis

YPAD overnight mid-log cultures were normalized to 7.10^6^ cells/ml and blocked in G1 with αfactor addition for two hours at 30°C. Dimethyl sulfoxide (DMSO) or camptothecin (CPT) were added directly to YPAD medium and incubated for 30min before release into S-phase. Samples were retrieved every 30min and directly mounted onto slides for analysis without fixation. For visualization of foci inside cells, images were recorded on a Zeiss AxioImager at 40X. Excitation of Rad52-YFP, Rfa1-CFP and Pus1-mCherry were respectively performed at 514, 439 and 587nm. Analysis of foci per cell was performed on the CellProfiler software with personalized pipeline (see Data Availability). Pus1-mCherry was used to mark the whole nucleus and count cells.

### Colony-forming viability assay

Cells grown for a time course like previously described were treated one hour with either DMSO or CPT before synchronous release still in the presence of drug. At each indicated time point, 300µL of cells at ∼1.10^7^ cells/mL were retrieved and diluted several times. The equivalent of 200 cells were spread on YPAD plates devoid of drug. DMSO-treated cultures were used as a spreading control, so the number of colonies that developed after CPT exposure was normalized to its corresponding DMSO plate. This value was then used to do a ratio between wild-type and *slx4Δ fun30Δ* survival rate.

### Chromatin immunoprecipitation

As previously mentioned, YPAD overnight mid-log cultures were normalized to 7.10^6^ cells/ml and blocked in G1 with αfactor addition for two hours at 30°C. Cells were then washed to remove the YPAD medium and resuspended in a permeabilizing medium (11) containing either DMSO or 50µM CPT still in the presence of αfactor for one more hour of incubation and treated with 1mM final Auxin Indole-3-acetic acid sodium salt (Merck, I5148) every hour for depletion of AID-tagged proteins. Cells were synchronously released into S-phase via pronase (50µG/mL) addition to the culture medium.

50 mL of 1.10^7^ cells were retrieved at indicated time points. For IP of Top1-ccs, no chemical crosslink was performed and cells were directly washed with cold TBS1X. For IP of RPA, cells were crosslinked for 15 min with 1% formaldehyde (Sigma F8775) at 25°C under agitation. Fixation was quenched by addition of 250 mM Glycine (Sigma G8898) for 5 min under agitation. Cells were washed two times with cold TBS1X (4°C). All dry pellets were frozen and conserved at −20°C until processed. Cell pellets were resuspended in lysis buffer (50 mM HEPES-KOH pH7.5, 140 mM NaCl, 1 mM EDTA, 1% Triton X-100, 0.1% Na-deoxycholate) supplemented with 1 mM PMSF and anti-protease (cOmplete Tablet, Roche, 505649001) and lysed by beads-beat method (MB400 U, Yasui Kikai, Osaka). Recovered lysate was either sonicated directly with a BioRuptor® Pico system (Diagenode) (6 cycles: 30s ON, 30s OFF) or completed to 3 mL with cold lysis buffer and sonicated with a Q500 sonicator (Qsonica) (12 cycles: 15 s ON, 45 s OFF, amplitude 50). Dynabeads were washed three times and resuspended in 1 mL of PBS, 0.5% BSA, 0.1% Tween and incubated with antibodies on a rotating wheel for six hours at 4°C. We used for each IP condition either 0,5µl of anti-RPA (Agrisera, AS07214) with 100µl Dynabeads Prot. A (DPA); or 2,5µL anti-FLAG (Sigma, F3165) with 80µL of DPA. Antibodies-coupled Dynabeads were washed three times with 1 mL of PBS, 0.5% BSA, 0.1% Tween and added to ∼1000µL of WCE overnight on a rotating wheel at 4°C. Samples of WCE were kept for the Input sample (5%), sonication control and for western-blotting (WB). After IP, beads were collected on a magnetic rack. A sample of the supernatant was collected for WB control (Flow-Through sample) and beads were washed on ice with cold solutions: two times with Lysis buffer (50 mM HEPES-KOH pH7.5, 140 mM NaCl, 1 mM EDTA, 1% Triton X-100, 0.1% Na-deoxycholate), twice with Lysis buffer + 0.36 M NaCl (50 mM HEPES-KOH pH7.5, 360 mM NaCl, 1 mM EDTA, 1% Triton X-100, 0.1% Na-deoxycholate), twice with Wash buffer (10 mM Tris-HCl pH8, 0.25 M LiCl, 0.5% IGEPAL, 1 mM EDTA, 0.1% Na-deoxycholate) and once with RT TE (10 mM Tris-HCl pH8, 1 mM EDTA). Antibodies were un-coupled from beads with 120µL of Elution Buffer (50 mM Tris-HCl pH8, 10 mM EDTA, 1% SDS) for 11 min at 65°C with occasional shaking. Eluates were collected and incubated with either 120µL of TE alone for non-crosslinked samples or 120 µL of TE, 0.1%SDS for de-crosslinking. Crosslinked samples were incubated at 65°C overnight while non-crosslinked samples directly followed the next steps. 190 µL of TE containing 60 mg RNase A (Sigma, R65-13) were added to the samples and incubated for 2 hours at 37°C. Proteins were digested by addition of 20µL of Proteinase K (Sigma, P6556) at 20 mg/ml and incubated for 2 hours at 37°C. 50µL of 5M LiCl were added to DNA before purification by two rounds of Phenol: Chloroform: Isoamyl Alcohol 25:24:1 (Sigma, P2069) extractions and precipitation by addition of 100mM Sodium Acetate (Sigma, S2889), 26 mg/ml of Glycogen (Roche, 10901393001) and 1mL of 100% ethanol overnight at −20°C. Samples were centrifuged for 45 min at 15.000 rpm at 4°C, washed with cold 70% ethanol and centrifuged again 5 min at 4°C. DNA pellets were dried and resuspended in H20 or TE before quantification and adjustment prior to qPCR reactions. qPCR reaction was performed with LightCycler480 (Roche). IP/Input ratios were calculated and qPCR results of each experiment were normalized on *NEGV* intergenic region (coordinates 469104-469177 on chromosome V), negative for Top1 and RPA binding and DNA replication in our experimental conditions.

## Results

### Role of the S-phase checkpoint in response to CPT-induced replication stress

Although CPT is well characterized as an inducer of replication-dependent DNA damage, it has long been assumed that acute CPT exposure does not significantly impair DNA replication kinetics in budding yeast (10, 35, 36). In a previous study, we challenged this assumption by demonstrating that increasing the cell permeability to CPT induced a delay in the progression through the S- and G2/M phases of the cell cycle compared to untreated controls (11). Since the response to DNA damage during DNA replication is coordinated by the S-phase checkpoint, we investigated whether the cell cycle delays observed in CPT-treated cells depend on this pathway.

To this end, we monitored cell cycle progression in wild-type and checkpoint mutant cells by flow cytometry following synchronization in G1 and release into a single cell cycle (**Figure 1A**). We analyzed mutants affecting key components of the checkpoint pathway, including the sensor kinase Mec1, the effector kinase Rad53, and the mediators Rad9 and Mrc1, which defines two distinct branches of Rad53 activation (27). As deletion of *MEC1* and *RAD53* is lethal, the viability of *mec1Δ* and *rad53Δ* mutants was rescued by deleting *SML1*, which encodes an inhibitor of the ribonucleotide reductase, allowing *de novo* dNTP synthesis during S phase (37). Because *rad53Δ sml1Δ* cells grow more slowly than wild-type cells due to the accumulation of free histones (38) (Figure 1C), and Rad53 checkpoint function is not required for dNTP pool maintenance in control conditions (39), we used the kinase-deficient mutant *rad53-11* (*G653E*), which retains normal growth and dNTP synthesis ability but is defective for checkpoint activation in response to DNA damage (40). Since Mrc1 is an integral component of the replisome and promotes DNA replication under normal conditions (41–43), we analyzed both *mrc1Δ* and the phosphorylation-deficient *mrc1-AQ* allele, which is specifically defective in checkpoint signaling.

**Figure 1.**
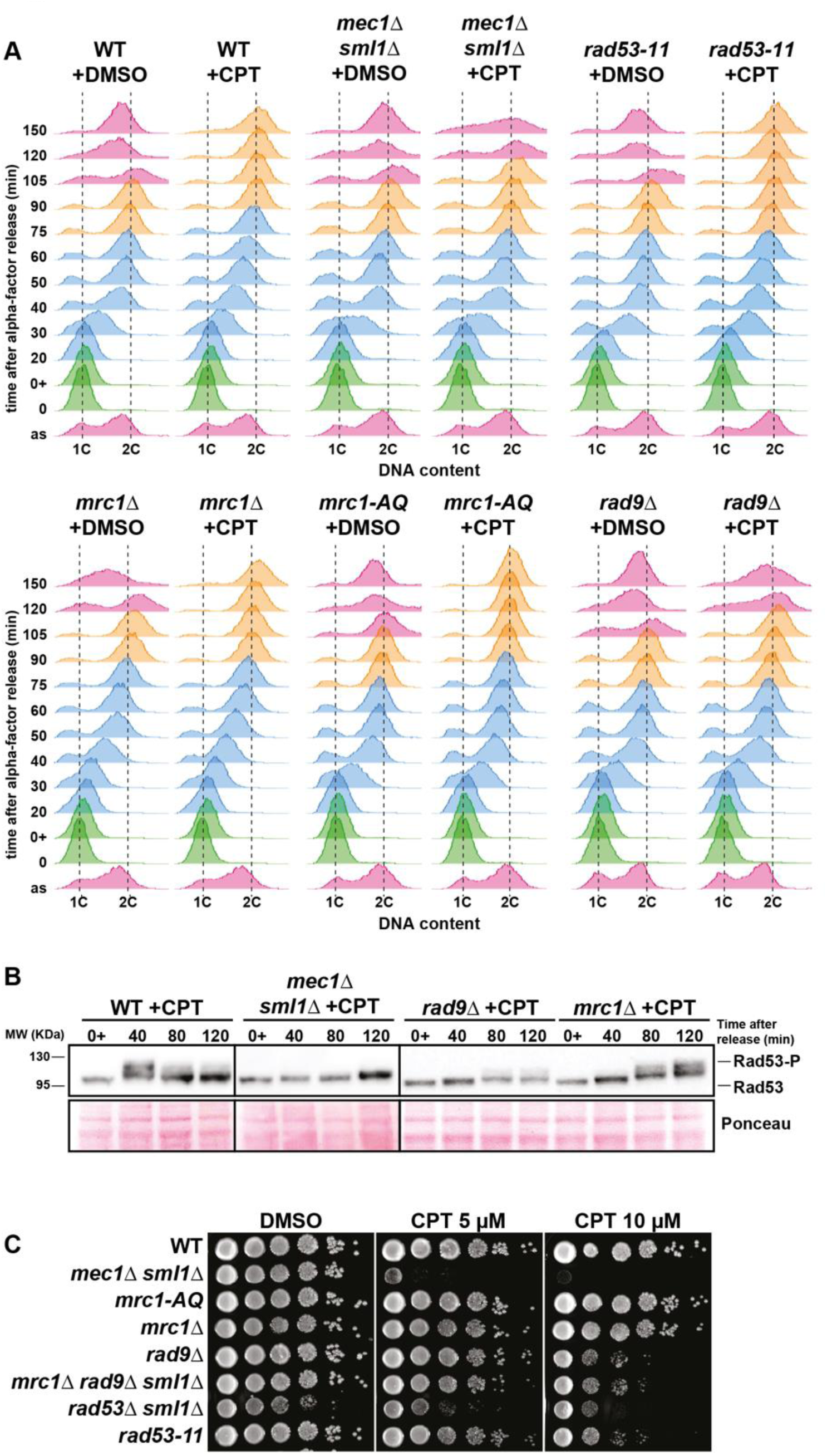
CPT-induced DNA damage activates DDC-dependent cell cycle delays important for cell viability. **A** - Analysis of DNA content (1C, 2C) by flow cytometry in a time course experiment. Cells were collected at the indicated time points after being blocked in G1 phase with α-factor (0), treated with 50µM CPT or DMSO for one hour (0+) in MPD +SDS medium and synchronously released into S phase still in presence of drug. Time spent in un-synchronized cell cycles (pink), G1 (green), S (blue) and G2/M (orange) are colored. See text for more details. Three biological replicates have been performed. **B** –Rad53 phosphorylation analyzed by western blot in wild-type and indicated mutants. Cells were blocked in G1 by using α-factor, treated with 100µM CPT for one hour (0+) in MPD +SDS medium and synchronously released into S-phase still in presence of drug. Cells were collected at the indicated time points and Rad53 was immunodetected with an antibody against total Rad53. Three biological replicates have been performed. **C** – CPT sensitivity assay of indicated genotypes. 10-fold serial dilutions of yeast cells were spotted on YPAD growth plates containing increasing doses of CPT and incubated for 2 days at 30°C. Three biological replicates have been performed.

**Supplementary Figure 1.**
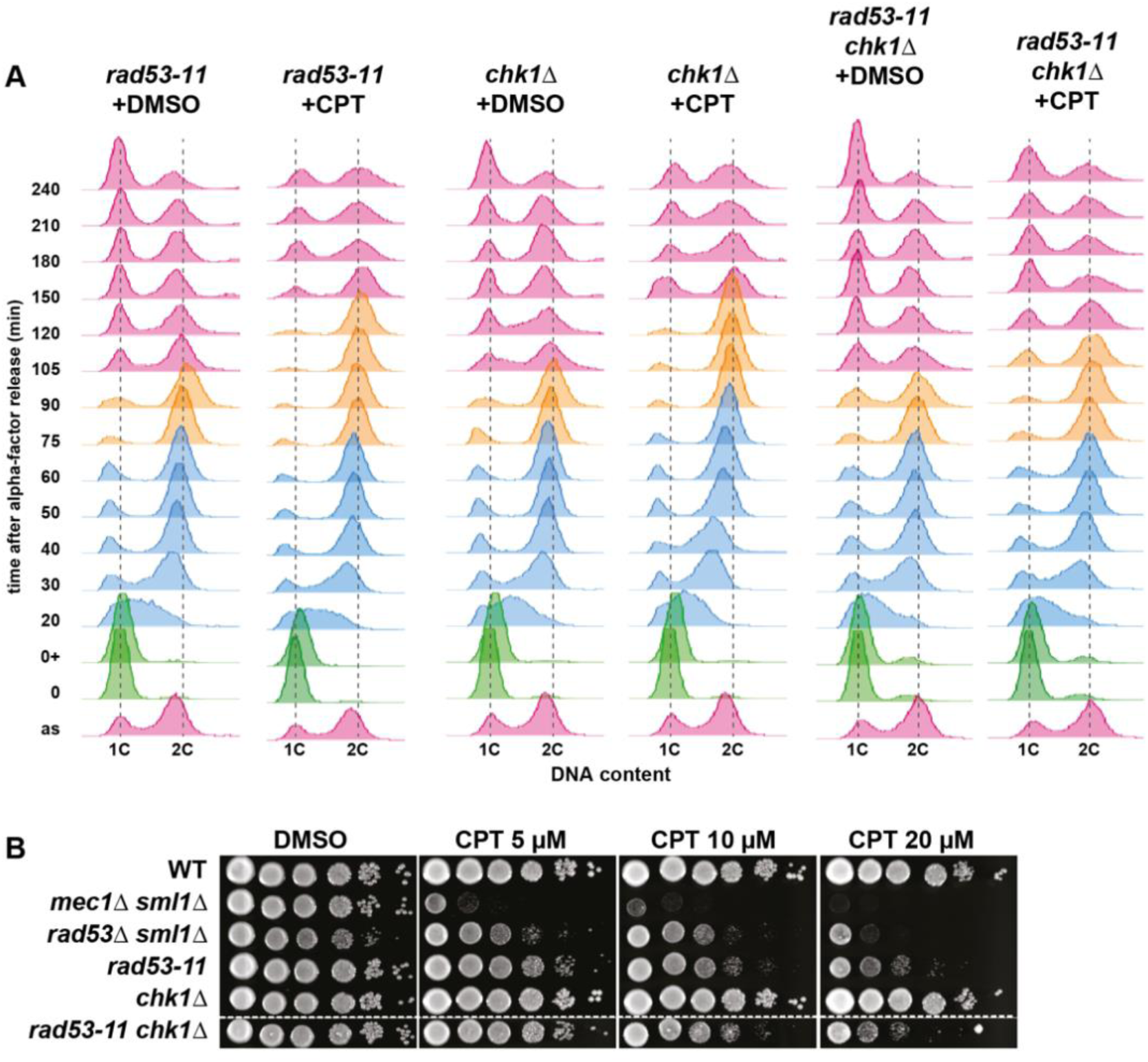
The G2/M cell cycle delay induced by CPT treatment is dependent on Rad53 and Chk1. **A** - Analysis of DNA content (1C, 2C) by flow cytometry in a time course experiment. Cells were collected at the indicated time points after being blocked in G1 phase with α-factor (0), treated with 50µM CPT or DMSO for one hour (0+) in MPD +SDS medium and synchronously released into S phase still in presence of drug. Time spent in un-synchronized cell cycles (pink), G1 (green), S (blue) and G2/M (orange) are colored. See text for more details. Three biological replicates have been performed. **B** – CPT sensitivity assay of indicated genotypes. 10-fold serial dilutions of yeast cells were spotted on YPAD growth plates containing increasing doses of CPT and incubated for 2 days at 30°C. Three biological replicates have been performed.

Asynchronous cell cultures were first arrested in G1 phase with α-factor mating pheromone (time point 0), then treated with either DMSO or CPT for one hour (time point 0^+^). Following treatment, cells were synchronously released into S phase in the presence of drug. Samples were collected at indicated time points, fixed and stained with propidium iodide, a DNA intercalant whose fluorescence intensity was measured by flow cytometry. The intensity ranged from one DNA content (1C) in G1 to two DNA content (2C) when cells reach G2/M. Dotted lines corresponding to the 1C and 2C peaks in the WT+DMSO condition were drawn and applied across all strains. Using this approach, the duration of S phase was defined as the interval until the bulk of cells reached the 2C dotted line, and the duration of G2/M phase was assessed by the subsequent decline of the 2C population. For clarity, cell cycle phases (asynchronous, G1 block, S and G2/M) were color coded (pink, green, blue and orange, respectively). Wild-type cells treated with DMSO entered S phase approximately 20 minutes after release from the G1 block and reached G2/M by 75 minutes. Consistent with previous observations (11), CPT treatment induced a clear delay in S-phase progression, with wild-type cells reaching G2/M only 90 minutes post release (**Figure 1A**). This CPT-induced delay was not detected in *mec1Δ sml1Δ* and *rad53-11* mutants, which displayed similar S-phase kinetics regardless of treatment with DMSO or CPT (**Figure 1A**). These results indicate that the Mec1-Rad53 pathway delays S-phase progression in response to CPT-induced DNA damage. *mrc1Δ* and *mrc1-AQ* cells behaved like wild-type cells, showing an S-phase delay when treated with CPT compared to the control conditions (+DMSO). In contrast, the *rad9Δ* mutant progressed through S phase with kinetics comparable to DMSO-treated controls, regardless of CPT exposure (**Figure 1A**). This suggests that the CPT-induced S-phase delay depends on the DDC branch of the S-phase checkpoint, involving Mec1, Rad9 and Rad53.

Furthermore, CPT-treated wild-type cells exhibited an extended G2/M phase relative to controls, consistent with checkpoint-mediated inhibition of anaphase entry (Clarke et al., 2001; Redon et al., 2003; Zhou et al., 2016). This delay was partially suppressed in *mec1Δ sml1Δ* and *rad9Δ* cells but persisted in the *rad53-11* mutant (**Figure 1A**). These results imply the existence of another checkpoint effector kinase activated by Mec1 and Rad9 and that acts redundantly with Rad53 to maintain the G2/M block. Such a function has been described for the kinase Chk1 (45). Indeed, deletion of *CHK1* in the *rad53-11* mutant partially relieved the CPT-induced G2/M delay (**Supplementary Figure 1A**).

The activation of S-phase checkpoints was then assessed by the phosphorylation status of Rad53 in wild-type and mutant cells, which is monitored as an electrophoretic mobility shift by classical Western blot (**Figure 1B**). Asynchronous cultures were blocked in G1 phase with α-factor, then treated with CPT for one hour and synchronously released into S-phase in the continued presence of the drug. Protein samples were collected at different time points spanning a full cell cycle. In contrast to previous reports (35, 36), we observed a weak but significant Rad53 phosphorylation upon S-phase entry at 40 minutes after release (10) followed by dephosphorylation at later time points (**Figure 1B**). No S-phase-specific Rad53 phosphorylation was detected in *mec1Δ sml1Δ* and *rad9Δ* mutants, while Rad53 phosphorylation occurred in *mrc1Δ* mutants, but was delayed. These results are consistent with the replication delays observed in these mutants in response to CPT (**Figure 1A**), confirming the role of the DDC branch (Mec1-Rad9-Rad53) of the S-phase checkpoint in slowing down DNA replication in response to CPT-induced DNA damage.

To investigate the functional significance of S-phase and G2/M delays, we assessed the CPT sensitivity of various checkpoint mutants (**Figure 1C**). Serial dilutions of cells were spotted on plates containing increasing concentrations of CPT, and colony formation was used as a readout of cellular fitness under CPT-induced replication stress. Wild-type, *mrc1Δ* and *mrc1-AQ* mutants displayed no detectable sensitivity to CPT. In contrast, *mec1Δ sml1Δ* cells were highly sensitive to CPT, even at the lowest dose tested (CPT 5 µM). Similar to *mec1Δ*, the *rad9Δ mrc1Δ* double mutant is lethal but can be suppressed by deleting *SML1*, enabling further analysis. Interestingly, *rad9Δ*, *rad9Δ mrc1Δ sml1Δ* and *rad53-11* mutants were resistant to 5 µM CPT, but displayed a similar increased sensitivity at 10 µM relative to control cells (**Figure 1C**). These findings support several conclusions: (*i*) The cell cycle delays induced by DDC (**Figure 1A,B**) contribute to cell survival in response to CPT. (*ii*) The comparable sensitivity of *rad9Δ and rad9Δ mrc1Δ* mutants suggests that Mrc1 cannot compensate for the loss of Rad9 for CPT resistance. *(iii*) The greater sensitivity of *mec1Δ sml1Δ* cells compared to *rad53-11* and *rad9Δ* mutants suggests that Mec1 has functions independent of the canonical checkpoint cascade in maintaining genome stability (46, 47). We also monitored the CPT sensitivity of *rad53-11 chk1Δ* mutants, which lack the G2/M delay (**Supplementary Figure 1A**). Surprisingly, deletion of *CHK1* did not reduce the viability of the *rad53-11* mutant to CPT exposure, indicating a more complex relationship between cell cycle delays and viability in response to CPT (**Supplementary Figure 1B**).

Collectively, our results show that the S-phase checkpoint, and more specifically the DDC branch, induces cell cycle delays that may be important for cell viability in response to CPT-induced DNA damage.

### Activation of the S-phase checkpoint is locally dampened by Slx4-Rtt107 and Fun30

To explain the weak phosphorylation of Rad53 in wild-type cells exposed to CPT (**Figure 1B**), we hypothesized that checkpoint activity could be dampened by Slx4-Rtt107 in response to this particular replication stress, as described earlier for MMS and HU (30, 32). To address this possibility, we analyzed Rad53 phosphorylation in CPT-treated *rtt107Δ* and *slx4Δ* mutants. Compared to the wild type, these mutants displayed pronounced Rad53 hyperphosphorylation upon entry into S phase (t=40 min) in the presence of CPT, which persisted at later time points (**Figure 2A,B**). Importantly, Rad53 hyperphosphorylation was abolished in *slx4Δ rad9Δ* double mutants, confirming that this hyperactivation requires Rad9.

**Figure 2.**
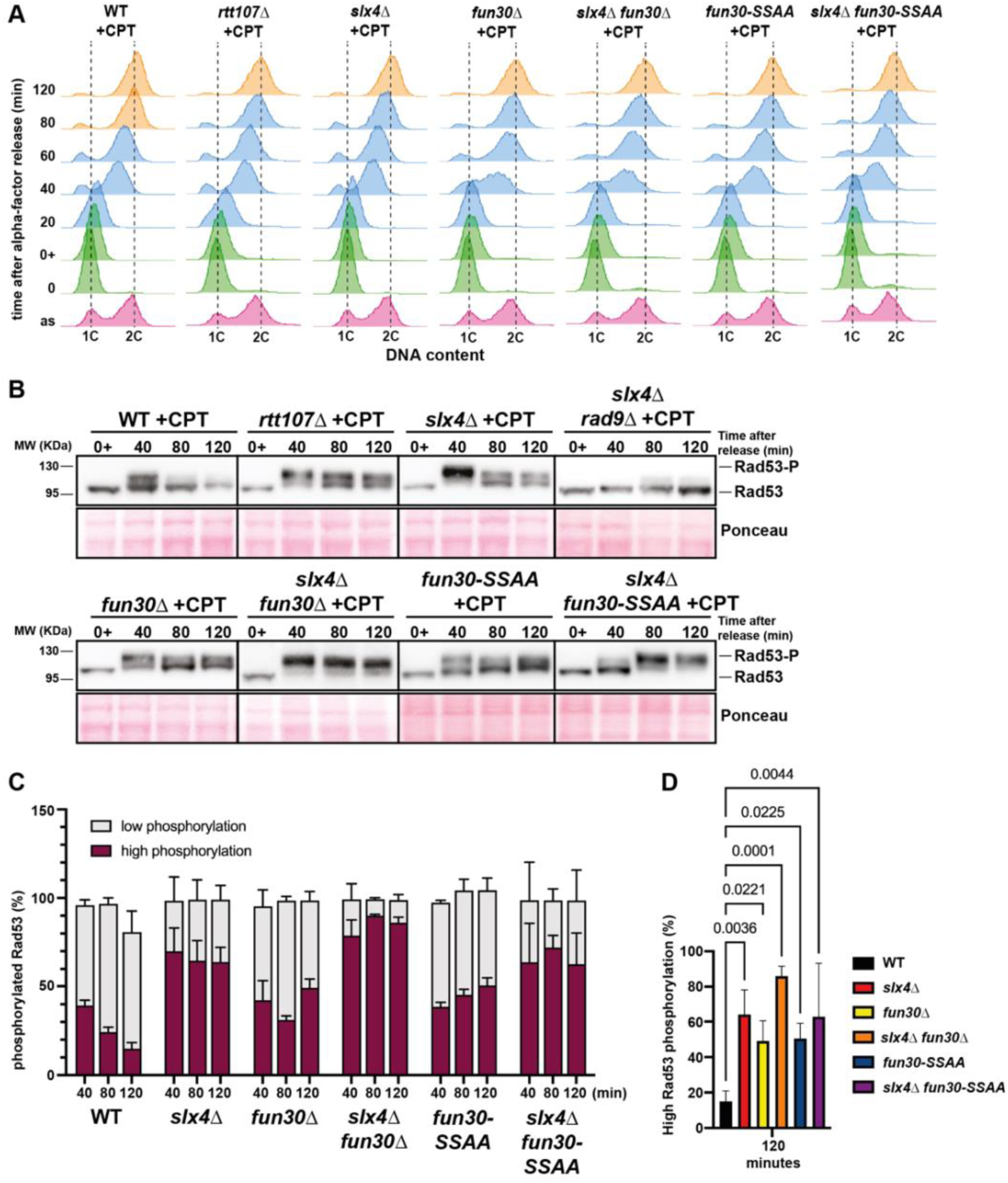
– Activation of the S-phase checkpoint in response to CPT is strongly dampened by the additive effect of Slx4-Rtt107 and Fun30 counteracting Rad9. **A** – Analysis of DNA content (1C, 2C) by flow cytometry of the experiment shown in panel B, performed as in Figure 1A. Profiles correspond to. Three biological replicates have been performed. **B** – Rad53 phosphorylation analyzed by western blot in wild-type and indicated mutants performed as in Figure 1B. Three to five biological replicates have been performed. **C** – Quantification of Rad53 phosphorylation showing the ratio of phosphorylated Rad53 (low and high)/total Rad53 observed by Western blot in indicated mutants. Data represent the mean ± SEM of three to five biological replicates. **D** – Statistical analysis of Rad53 high phosphorylation at 120 minutes after release into S phase. Data represent the mean ± SEM of three to five biological replicates. P-values of pairwise comparisons to the wild-type tested by one-way ANOVA are indicated.

The chromatin remodeler Fun30 has also been shown to counteract Rad9 functions in response to DNA damage (48–51). Notably, Fun30 loss resulted in prolonged Rad53 hyperphosphorylation in response to irreparable DSB (49). Since Fun30 and Rad9 interact with the same domain of Dpb11, it could compete with Rtt107-Slx4 to displace Rad9 from DNA damage sites (52). Indeed, Fun30 limits the activation of Rad9-dependent DDC in cells exposed to MMS and CPT (50). Consistent with this model, we found that the absence of Fun30 resulted in prolonged Rad53 hyperphosphorylation during the time course (**Figure 2A,B**). Moreover, *fun30Δ* cells expressing the *fun30-SSAA* allele, which specifically abolishes the interaction between Fun30 and Dpb11 (52), also exhibited checkpoint hyperactivation, suggesting that Fun30’s physical interaction with Dpb11 is required to displace Rad9 and dampen checkpoint signaling (**Figure 2A,B**). This effect was additive with the absence of Slx4, as shown by the higher percentage of hyperphosphorylated Rad53 in *slx4Δ fun30Δ* and *slx4Δ fun30-SSAA* compared to *slx4Δ*, *fun30Δ* or *fun30-SSAA* single mutants (**Figure 2C,D**). Together, these results suggest that Slx4-Rtt107 and Fun30 function independently to attenuate the DNA damage checkpoint by limiting Rad9’s binding to Dpb11, thereby fine-tuning the checkpoint response to CPT-induced replication stress.

We next used flow cytometry to examine how prolonged checkpoint activation affects cell cycle progression in *slx4Δ fun30Δ* cells exposed to CPT (**Figure 3A**). Under control conditions (+DMSO), *slx4Δ fun30Δ* cells progressed through the cell cycle similarly to wild type cells, indicating that Slx4 and Fun30 are not required for cell cycle progression in the absence of replication stress (**Figure 3A**). However, following CPT treatment, *slx4Δ fun30Δ* cells showed a more pronounced S phase delay compared to wild type, consistent with sustained checkpoint activation in the absence of these checkpoint dampening factors.

**Figure 3.**
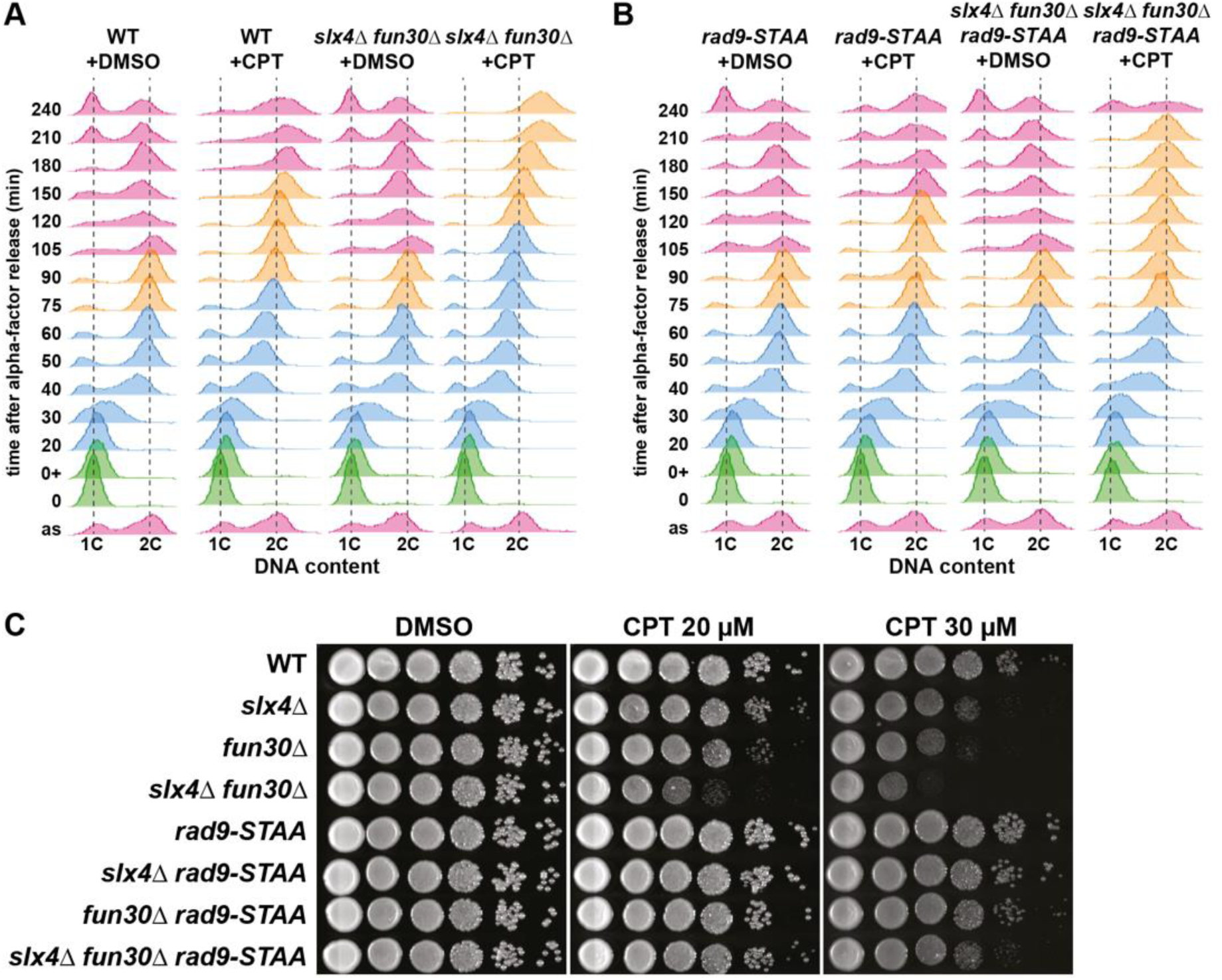
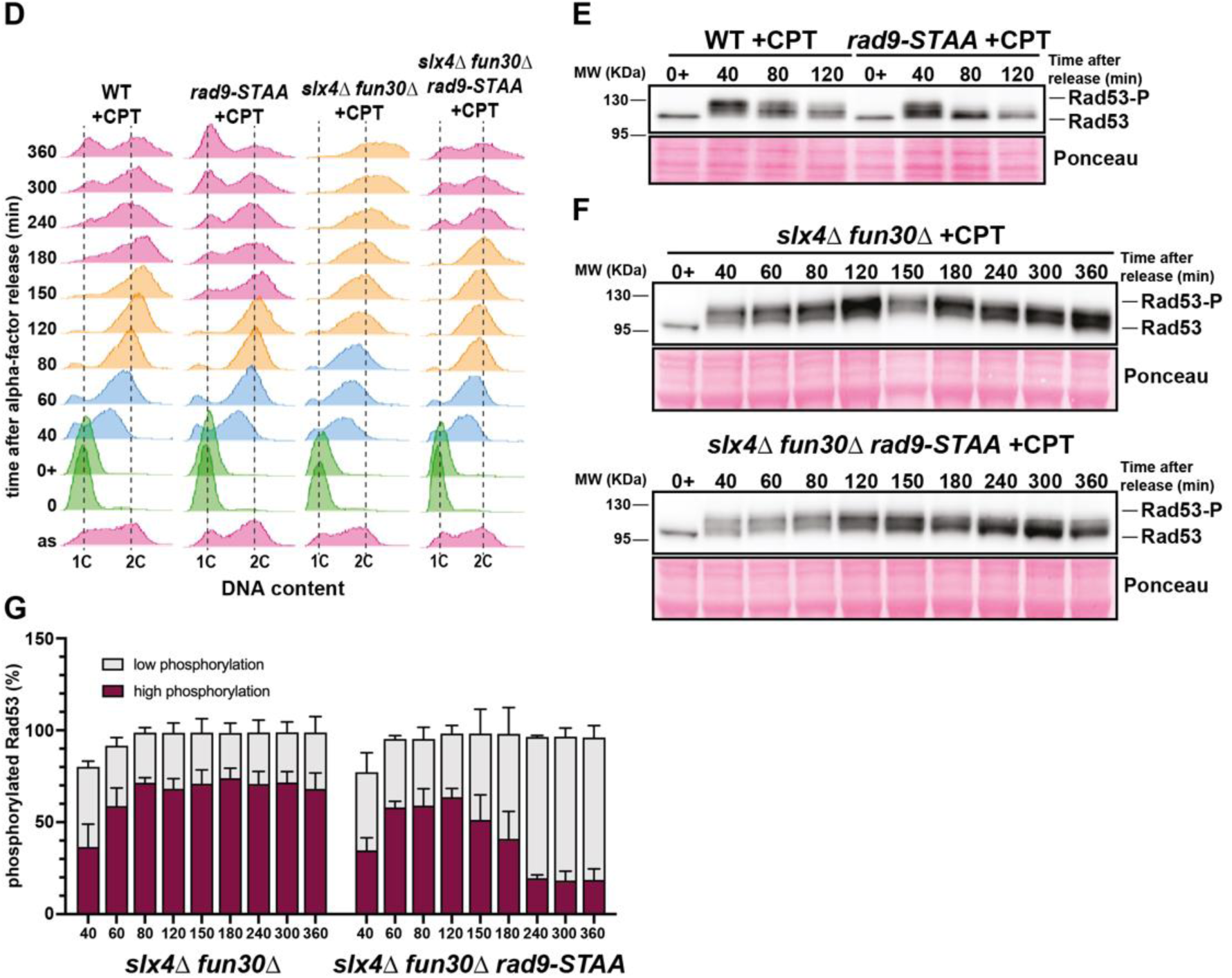
–Loss of Rad9 binding to Dpb11 counterbalances the checkpoint hyperactivation caused by the absence of Slx4 and Fun30. **A, B** – Analysis of DNA content (1C, 2C) by flow cytometry in a time course experiment performed as in Figure 1A. Three biological replicates have been performed. **C** – CPT sensitivity assay of indicated genotypes. 10-fold serial dilutions of yeast cells were spotted on YPAD growth plates containing increasing doses of CPT and incubated for 2 days at 30°C. Three biological replicates have been performed. **D** - Analysis of DNA content (1C, 2C) by flow cytometry of the experiments shown in panels E and F. **E, F** – Rad53 phosphorylation analyzed by western blot in wild-type and indicated mutants performed as in Figure 1B. Three biological replicates have been performed. **G** – Quantification of Rad53 phosphorylation showing the ratio of phosphorylated Rad53 (low and high)/total Rad53 observed by Western blot in indicated mutants. Data represent the mean ± SEM of three biological replicates.

To further characterize the DNA synthesis delay observed in *slx4Δ fun30Δ* cells, we analyzed the migration of full-length chromosomes using pulsed-field gel electrophoresis (PFGE), which separates of individual linear chromosomes based on size. During S phase, chromosomes accumulate replication intermediates - referred to here as joint molecules (JMs) - that fail to enter the gel and remain trapped in the wells. Ethidium bromide staining revealed that chromosome bands detectable in G1-arrested cells disappeared during S phase (40-60 minutes after release) and reappeared in G2 phase (t=80 min) in wild-type cells treated with CPT (**Supplementary Figure 3A**). In *slx4Δ fun30Δ* cells, chromosome disappeared from the gel at the same time as control cells, but completed DNA synthesis with a 20-minute delay (t=100 min) compared to the wild-type. This delay in chromosome migration was confirmed by Southern blot analysis of chromosome VI, which showed a strong accumulation of JMs in the well correlating with a loss of chromosome VI signal in the gel (**Supplementary Figure 3A, bottom panels**). These results are consistent with the S-phase delay observed by flow cytometry in the *slx4Δ fun30Δ* double mutant (**Figure 3A**). Moreover, flow cytometry revealed that these cells remained blocked in G2/M for an extended period (up to 240 minutes post release), whereas wild-type cells completed the cell cycle and initiated a new one by 180 minutes (**Figure 3A**). Together, these findings indicate that *slx4Δ* and *fun30Δ* mutants exhibit prolonged delays in both S and G2/M, reflecting sustained activation of the S phase checkpoint in response to CPT-induced DNA damage.

Finally, we assessed the viability of *slx4Δ* and *fun30Δ* mutants under chronic CPT exposure using spot assays on growth plates (**Figure 3C**). Both *slx4Δ* and *fun30Δ* single mutants showed reduced viability at high CPT concentrations compared to wild type cells. Notably, the CPT sensitivity of these mutants was additive, as the *slx4Δ fun30Δ* double mutant displayed a greater loss of viability than either single mutant alone. The *fun30-SSAA* mutant, which disrupts Fun30’s interaction with Dpb11, conferred intermediate CPT sensitivity compared to *fun30Δ*, but still showed additive effects when combined with *slx4Δ* (**Supplementary Figure 3B**). Taken together, our results show that Slx4-Rtt107 and Fun30, through their interaction with Dpb11, play critical roles in attenuating S-phase checkpoint signaling in response to CPT, and that this dampening is important for the control of cell cycle progression and viability under replication stress. At the same time, our data show that the activation of the S-phase checkpoint is equally important for the control of cell cycle progression and viability (**Figure 1 and Supplementary Figure 1**). Thus, our results highlight the need for a finely tuned DNA damage response to ensure optimal survival in response to DNA damage.

### Loss of Slx4 and Fun30 repair functions are not the primary causes of checkpoint hyperactivation

Since Slx4 and Fun30 have been primarily described as DNA repair factors (48, 49, 52–57), one could argue that their absence could impair DNA repair and thereby indirectly impede checkpoint deactivation. Slx4 acts as a platform for recruiting structure-selective endonucleases such as Mus81, Rad1 and Slx1 to resolve various DNA repair intermediates. However, deletion of these nucleases did not result in a checkpoint hyperactivation (**Supplementary Figure 3C**), suggesting that Slx4’s checkpoint dampening function is independent of its nuclease-recruitment role.

Fun30 has been shown to promote DSB end resection through its chromatin remodeling activity. To determine whether this activity contributes to checkpoint dampening, we monitored checkpoint activation in the ATPase-dead *fun30-K603R* mutant (57). Expression of the *fun30-K603R* allele in *fun30Δ* cells suppressed the prolonged Rad53 hyperphosphorylation observed in the deletion mutant at late time points (**Supplementary Figure 3D**). However, the *fun30-K603R* allele did not rescue the increased CPT sensitivity of *fun30Δ* cells (**Supplementary Figure 3E**), in agreement with previous observations (57).

We further expressed the *fun30-K603R-SSAA* allele, which lacks both the ATPase activity and the ability to bind Dpb11, in *fun30Δ* cells. In this background, Rad53 remained hyperphosphorylated throughout the time course (**Supplementary Figure 3D**). These results indicate that, in contrast to its ability to bind Dpb11, Fun30 chromatin remodeling activity is not required for checkpoint dampening. Thus, the role of Fun30 in checkpoint dampening is functionally distinct from its involvement in DNA repair.

To restore the unbalanced checkpoint response observed in *slx4Δ fun30Δ* mutant, we used the *rad9-STAA* allele, which prevents Rad9 binding to Dpb11. Despite this disruption, Rad9 can still be recruited to DNA damage sites, presumably via alternative binding partners, and retains the ability to activate the DDC in response to phleomycin-induced DNA breaks (58). Interestingly, *rad9-STAA* cells exhibited no increased sensitivity to CPT relative to wild-type cells (**Figure 3C**), indicating that the Rad9-Dpb11 interaction is dispensable for resistance to CPT-induced DNA damage. Strikingly, *rad9-STAA* almost completely suppressed the CPT sensitivity of *slx4Δ* and *fun30Δ* single mutants. *rad9-STAA* only partially suppressed the sensitivity of the *slx4Δ fun30Δ* double mutant, but the suppression was complete for the *slx4Δ fun30-SSAA* mutant (**Supplementary Figure 3B**), suggesting that the Rad9-Dpb11 interaction is a key contributor to the checkpoint hyperactivation phenotype. Flow cytometry analysis further showed that *rad9-STAA* suppressed both S and G2/M delays induced by CPT in wild-type and *slx4Δ fun30Δ* cells, although *slx4Δ fun30Δ rad9-STAA* cells were able to escape the G2/M block only 240 minutes after release (**Figure 3B**). The suppression of the S phase delay was also supported by PFGE experiments (**Supplementary Figure 3A**).

Consistent with these phenotypes, Rad53 phosphorylation kinetics were also altered in the *rad9-STAA* background. At 40 min post-release, Rad53 was phosphorylated in *rad9-STAA*, albeit to a lesser extent than in wild-type cells, and this phosphorylation declined over time (**Figure 3E**). In contrast, Rad53 remained hyperphosphorylated up to 360 minutes after release in the *slx4Δ fun30Δ* mutant, correlating with its sustained G2/M arrest (**Figure 3D**). In *slx4Δ fun30Δ rad9 STAA* cells, Rad53 phosphorylation was markedly reduced and slowly declined to low levels at 240 minutes, which coincided with cell cycle re-entry (**Figure 3D,F,G**). These findings show that loss of Rad9-Dpb11 interaction in the *rad9-STAA* mutant is sufficient to alleviate persistent checkpoint signaling and cell cycle arrest in *slx4Δ fun30Δ* cells, without restoring Slx4 and Fun30 repair functions.

Collectively, our results suggest that the checkpoint hyperactivation observed in the absence of Slx4 and Fun30 results from excessive Rad9-dependent activation of Rad53 at sites of DNA damage. This leads to a sustained G2/M arrest, and ultimately reduced viability upon CPT exposure. Thus, local checkpoint dampening by Slx4 and Fun30 is essential to permit cell cycle restart and survival in response to CPT-induced replication stress.

**Supplementary Figure 3.**
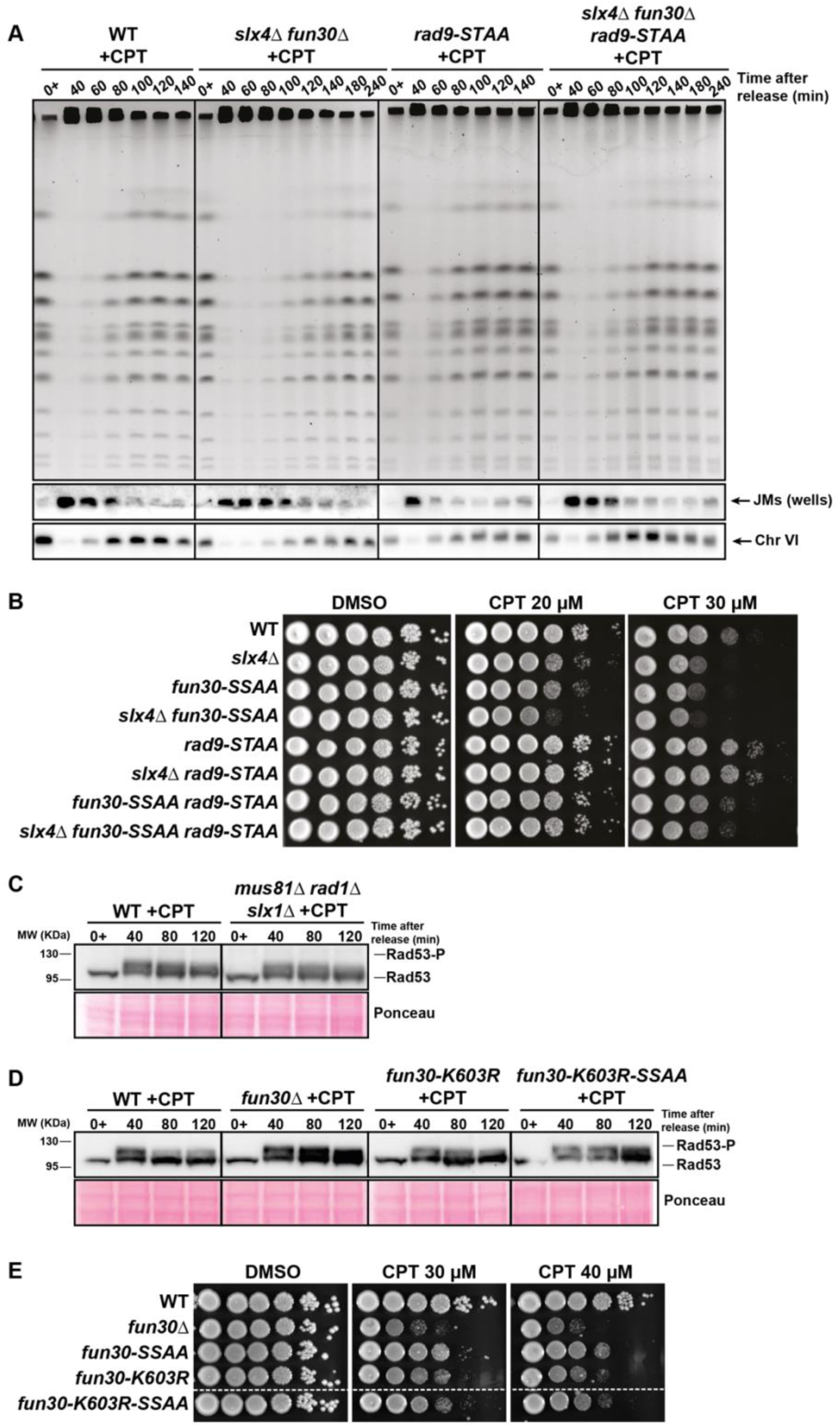
Loss of Slx4 and Fun30 repair functions are not the primary causes of checkpoint hyperactivation. **A** – Analysis of DNA replication by pulsed-field gel electrophoresis (PFGE) in indicated genotypes. Cells were treated as in Figure 1B and collected at the indicated time points. DNA extracted in agarose plugs was analyzed by PFGE. Upper panel: agarose gel stained with ethidium bromide. Lower panels: Southern blot using a probe specific to chromosome VI. JMs, joint molecules accumulated in the gel wells. Three biological replicates have been performed. **B, E** – CPT sensitivity assay of indicated genotypes. 10-fold serial dilutions of yeast cells were spotted on YPAD growth plates containing increasing doses of CPT and incubated for 2 days at 30°C. Three biological replicates have been performed. **C, D** – Rad53 phosphorylation analyzed by western blot in wild-type and indicated mutants. Cells were blocked in G1 by using α-factor, treated with 100µM CPT for one hour (0+) in MPD +SDS medium and synchronously released into S-phase still in presence of drug. Cells were collected at the indicated time points and Rad53 was immunodetected with an antibody against total Rad53. Three biological replicates have been performed.

### Hyperactivation of the S-phase checkpoint is deleterious for DNA repair

Although the hypersensitivity of *slx4Δ fun30Δ* cells to CPT might be attributed to prolonged G2/M arrest, it remains unclear whether checkpoint hyperactivation also disrupts DNA repair, independently of the canonical repair functions of Slx4 and Fun30. To address this possibility, we tested whether *slx4Δ fun30Δ* cells could exit G2/M arrest following CPT removal. In this recovery experiment, wild-type and *slx4Δ fun30Δ* cells were exposed to CPT for 120 minutes after release from α-factor and CPT was washed out. Cells were then allowed to recover overnight and the Rad53 phosphorylation was assessed by Western blot. Both strains showed Rad53 dephosphorylation upon recovery (**Figure 4A**), indicating that checkpoint signaling was turned off. However, colony formation assays revealed a loss of viability in the *slx4Δ fun30Δ* mutant compared to wild-type cells (**Figure 4B**). Notably, a 50% reduction in viability was observed when cells were exposed to CPT for at least 120 minutes, but not for 60 minutes, before drug removal. These results suggest that sustained checkpoint activation has a detrimental impact on cell survival.

**Figure 4.**
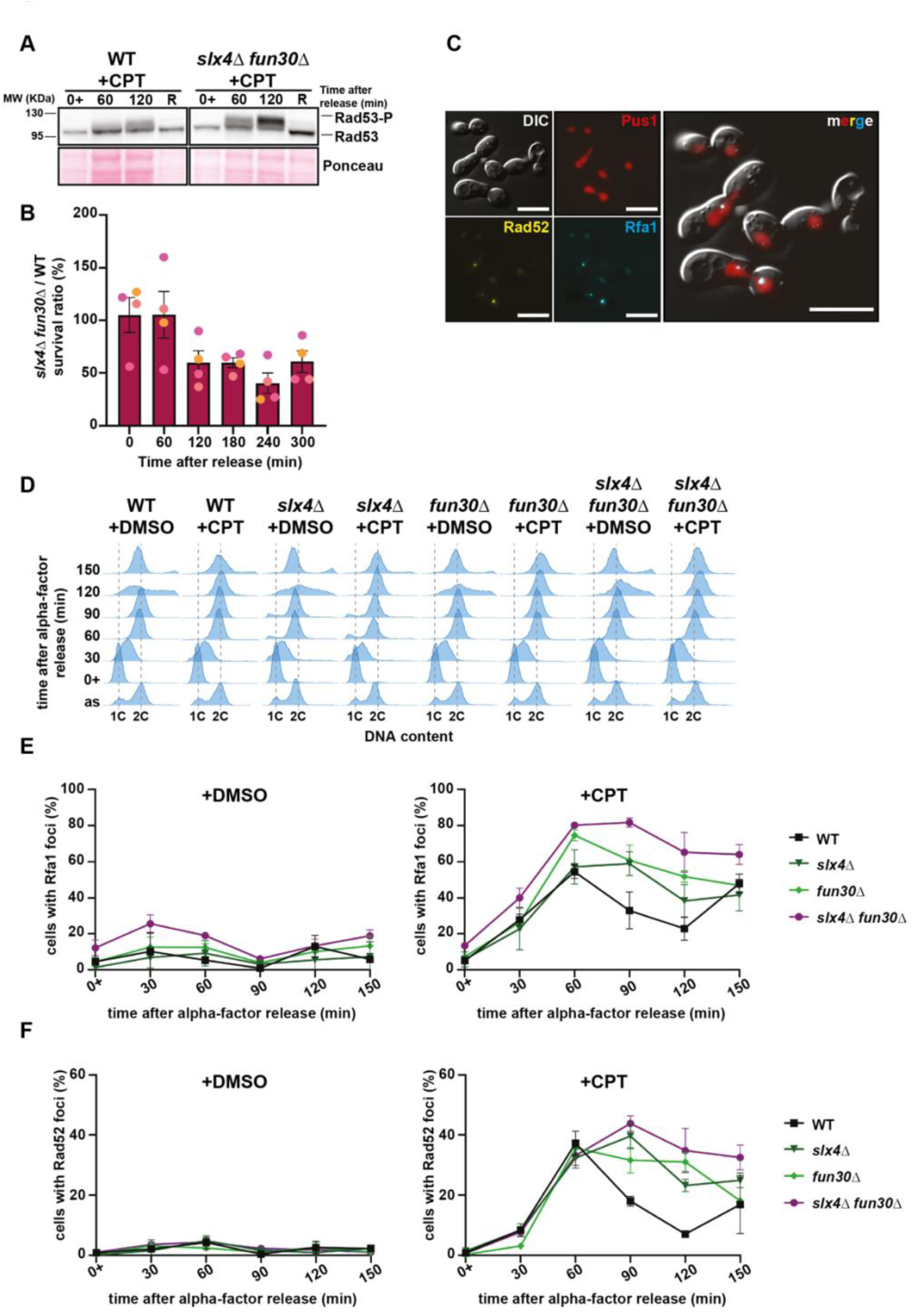
Checkpoint hyperactivation affects Homologous Recombination in response to CPT. **A** –Rad53 phosphorylation analyzed by western blot in wild-type and *slx4Δ fun30Δ* mutants. Cells were blocked in G1 by using α-factor, treated with 100µM CPT for one hour (0+) in MPD +SDS medium and synchronously released into S-phase still in presence of drug. 120 minutes after release, CPT was washed out and cells were put back to growth in YPAD medium to recover overnight (R). Three biological replicates have been performed. **B** – Colony-forming unit assay showing the ratio of survivor colonies in *slx4Δ fun30Δ*/wild-type cells. Cells were blocked in G1 by using α-factor, treated with 100µM CPT or DMSO for one hour (0+) in MPD +SDS medium and synchronously released into S-phase still in presence of drug. At each indicated time point after release, cells were washed three times and plated onto drug-free medium. Number of colonies previously exposed to CPT was normalized to that of colonies exposed to DMSO before calculating the mutant/wild-type ratio. Mean values ± SEM are plotted. Four biological replicates have been performed. **C** – Analysis of nuclear Rfa1-CFP and Rad52-YFP foci formation. An illustrative image of the experimental setup is shown. DIC, differential interference contrast; blue, Rfa1-CFP; yellow, Rad52-YFP; red, mCHERRY-Pus1 (nuclear compartment marker). Scale bar = 5 μm. **D** - Analysis of DNA content (1C, 2C) by flow cytometry of the experiments shown in panels E and F. **E, F** - Kinetic analysis of Rfa1-CFP (RPA) or Rad52-YFP foci formation in indicated genotypes. Cells were blocked in G1 by using α-factor, treated with 50µM CPT or DMSO for 30 minutes (0+) in YPAD medium and synchronously released into S-phase still in presence of drug. Cells were collected and imaged by fluorescence microscopy at the indicated time points. Quantification of foci was made with Cell Profiler software. Mean values ± SEM are plotted. Three to four biological replicates have been performed.

**Supplementary Figure 4.**
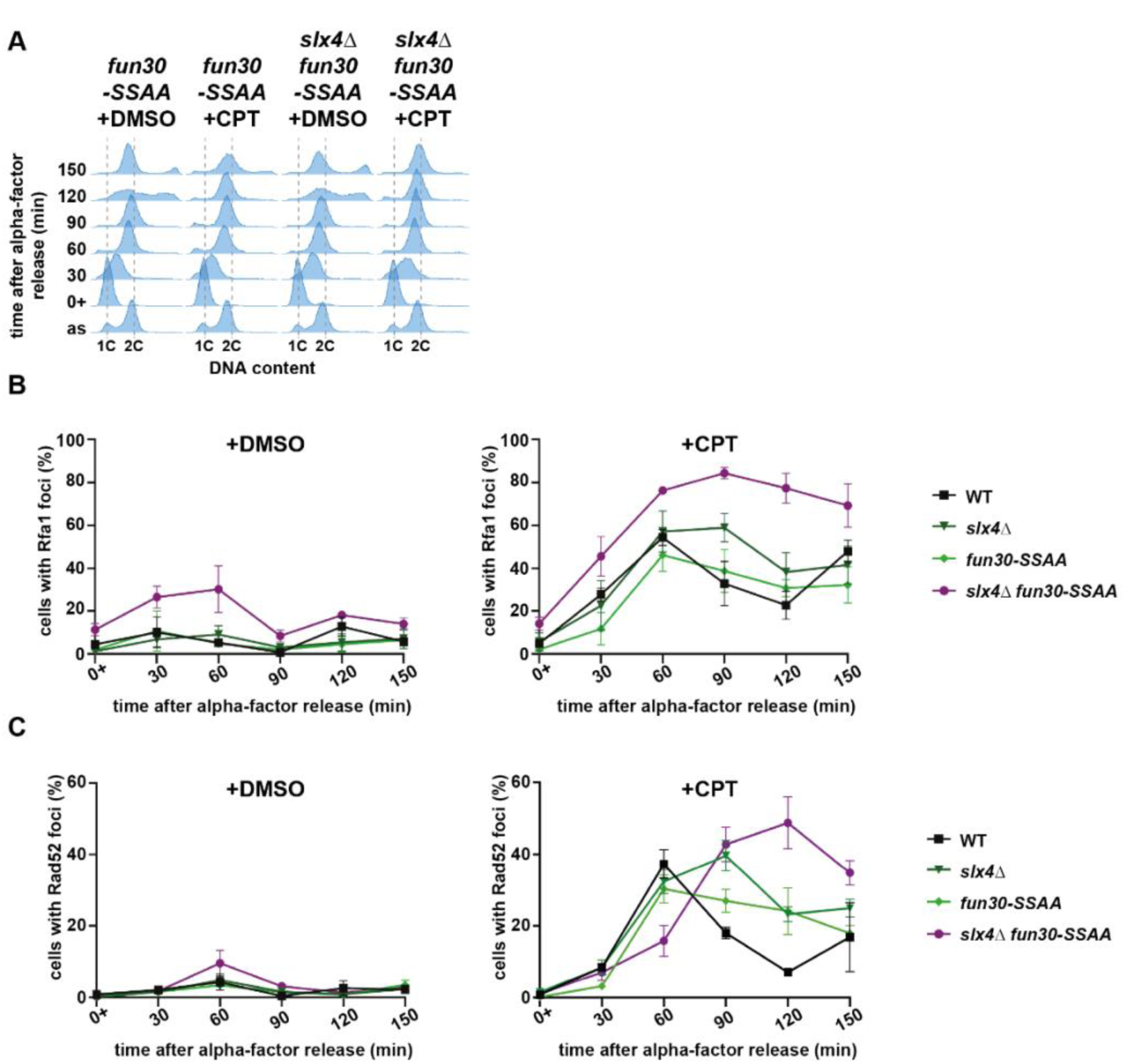
Homologous recombination is also affected in CPT-treated *slx4*Δ *fun30-SSAA* cells. **A** - Analysis of DNA content (1C, 2C) by flow cytometry of the experiments shown in panels B and C. **B, C** - Kinetic analysis of Rfa1-CFP (RPA) or Rad52-YFP foci formation in indicated genotypes. Cells were blocked in G1 by using α-factor, treated with 50µM CPT or DMSO for 30 minutes (0+) in YPAD medium and synchronously released into S-phase still in presence of drug. Cells were collected and imaged by fluorescence microscopy at the indicated time points. Quantification of foci was made with Cell Profiler software. Mean values ± SEM are plotted. Three to four biological replicates have been performed.

Sustained checkpoint signaling has been proposed to impair DNA repair (59–61). We previously reported that HR factors and the Mus81 endonuclease are paramount for the completion of DNA replication in cells exposed to CPT (11). Specifically, we proposed that HR factors protect reversed forks until they merge with a converging fork, forming an intermediate that is resolved by Mus81 during G2/M. To determine whether HR is affected by checkpoint hyperactivation in *slx4Δ fun30Δ* mutants exposed to CPT, we monitored the dynamics of HR protein foci in live cells. To this end, we engineered Rfa1 (RPA subunit) and Rad52 fluorescent tagged proteins and followed foci formation over time (**Figure 4C-F**). Cells were arrested in G1, treated with either DMSO or CPT, and then released synchronously into S phase, in the continued presence of the drug. Live cells were collected at 30-minute intervals, imaged by fluorescence microscopy, and the proportion of cells with at least one focus was quantified using an automated CellProfiler pipeline.

Wild-type cells treated with DMSO displayed low levels of Rfa1-CFP foci, with no more than 15% of cells showing foci during S phase, and fewer than 5% exhibiting Rad52-YFP foci. CPT treatment, however, induced a three-fold increase in the number of cells with Rfa1-CFP and Rad52-YFP foci, peaking at 60 minutes during S phase (**Figure 4E,F**). This was followed by a decrease at 90 and 120 minutes, as cells progressed through G2/M, likely reflecting the resolution of HR intermediates. The *slx4Δ fun30Δ* mutant showed low levels of Rfa1-CFP and Rad52-YFP foci in the absence of CPT, comparable to wild-type levels. Upon CPT exposure, *slx4Δ fun30Δ* cells showed a similar increase in S-phase foci compared to the wild type. However, Rfa1-CFP and Rad52-YFP foci persisted or accumulated over time (**Figure 4E,F**). Single *slx4Δ*, *fun30Δ*, and *fun30 SSAA* mutants also showed persistent foci, with levels intermediate between wild type and double mutants, consistent with the additive functions of Slx4 and Fun30 in checkpoint dampening (**Figure 4, Supplementary Figure 4**). Collectively, these results indicate that checkpoint hyperactivation interferes with the homologous recombination process in response to CPT exposure.

### Checkpoint hyperactivation does not prevent Mus81-Mms4 activation

Previous studies have suggested that the S-phase checkpoint could restrict the activation of the Mus81 endonuclease during HR-mediated DNA repair. In particular, premature activation of Mus81 has been observed in S-phase checkpoint mutants, inducing a premature processing of recombination intermediates and the generation of chromosome translocations (62). Conversely, it has been proposed that persistent checkpoint activity in the *slx4Δ pph3Δ* mutant could prevent the resolution of HR intermediates by Mus81 and recovery from MMS-induced DNA damage (61). In the context of CPT treatment, *mus81Δ* mutants also display persistent Rad52 foci, consistent with unresolved recombination intermediates (11). To determine whether the sensitivity of *slx4Δ fun30Δ* mutants to CPT reflects the inability of Mus81 to resolve CPT-induced HR intermediates, we analyzed genetic interactions between these mutants. As observed previously, *mus81Δ* cells were hypersensitive to low concentration of CPT (5 µM; **Figure 5A**). The combination of *mus81Δ* and *slx4Δ* did not show a clear additivity. However, CPT sensitivity was significantly increased in both *mus81Δ fun30Δ* and *mus81Δ slx4Δ fun30Δ* mutants. This additivity was also observed in the *mus81Δ slx4Δ fun30 SSAA* mutant (**Supplementary Figure 5A**), suggesting that checkpoint dampening and Mus81-mediated resolution function through distinct pathways. To further explore potential links between impaired Mus81 function and the *slx4Δ fun30Δ* phenotype, we introduced the *YEN1-ON* allele, which leads to premature activation of the Yen1 nuclease and can partially compensate for the loss of Mus81 (63). *YEN1-ON* slightly suppressed the CPT sensitivity of *slx4Δ fun30Δ* cells, but this increased resistance was already observed in the *YEN1-ON* single mutant compared to wild type (**Supplementary Figure 5B**). We conclude from these experiments that the loss of checkpoint dampening does not phenocopy the loss of Mus81.

**Figure 5.**
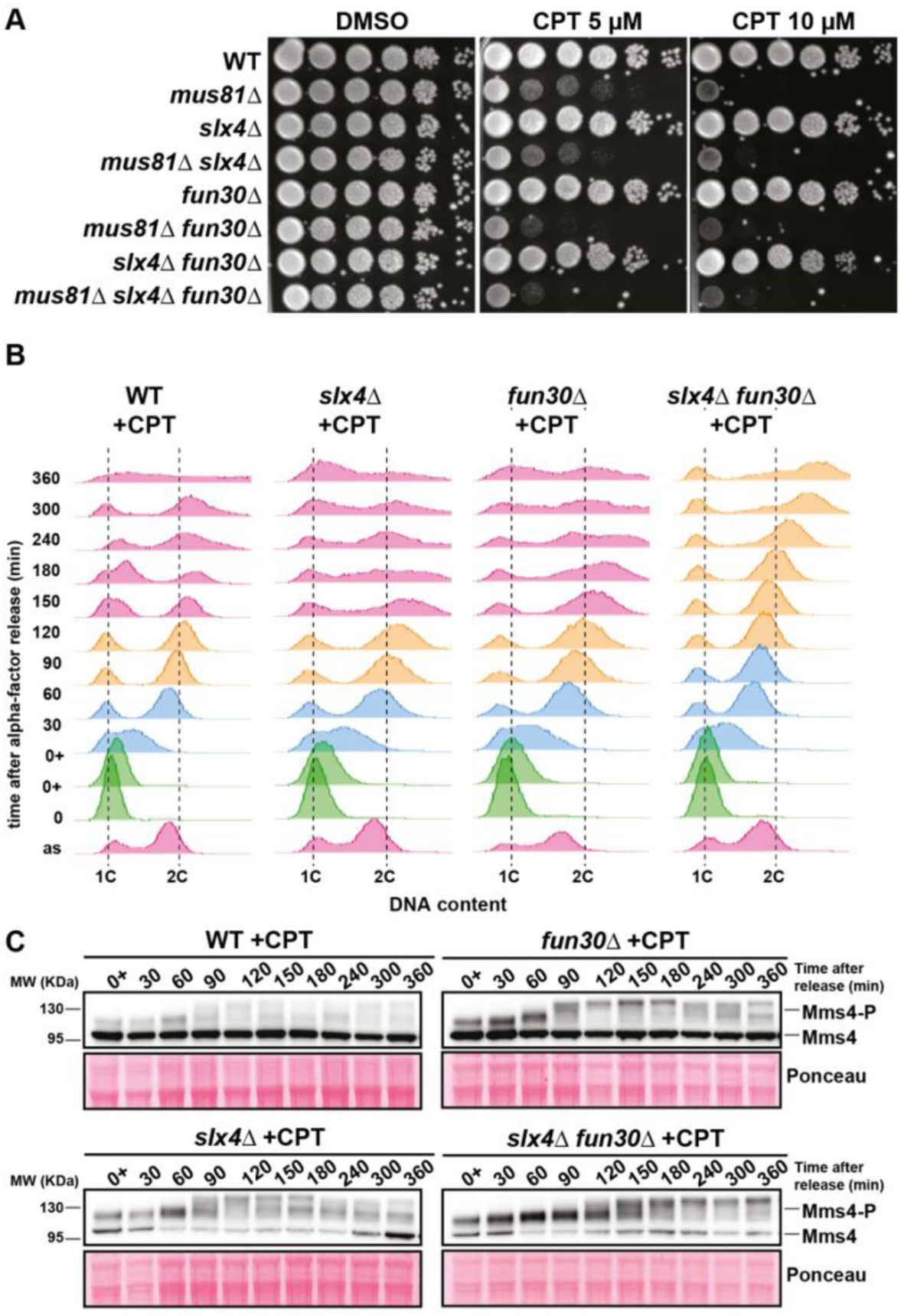
The absence of checkpoint dampening does not prevent the activation of Mus81-Mms4 in response to CPT. **A** – CPT sensitivity assay of indicated genotypes. 10-fold serial dilutions of yeast cells were spotted on YPAD growth plates containing increasing doses of CPT and incubated for 2 days at 30°C. Three biological replicates have been performed. **B** - Analysis of DNA content (1C, 2C) by flow cytometry of the experiments shown in panel C. **C** - Mms4 phosphorylation analyzed by western blot in wild-type and indicated mutants. Cells were blocked in G1 by using α-factor, treated with 100µM CPT for one hour (0+) in MPD +SDS medium and synchronously released into S-phase still in presence of drug. Cells were collected at the indicated time points and Mms4 was immunodetected with an anti-Flag antibody. Three biological replicates have been performed.

**Supplementary Figure 5.**
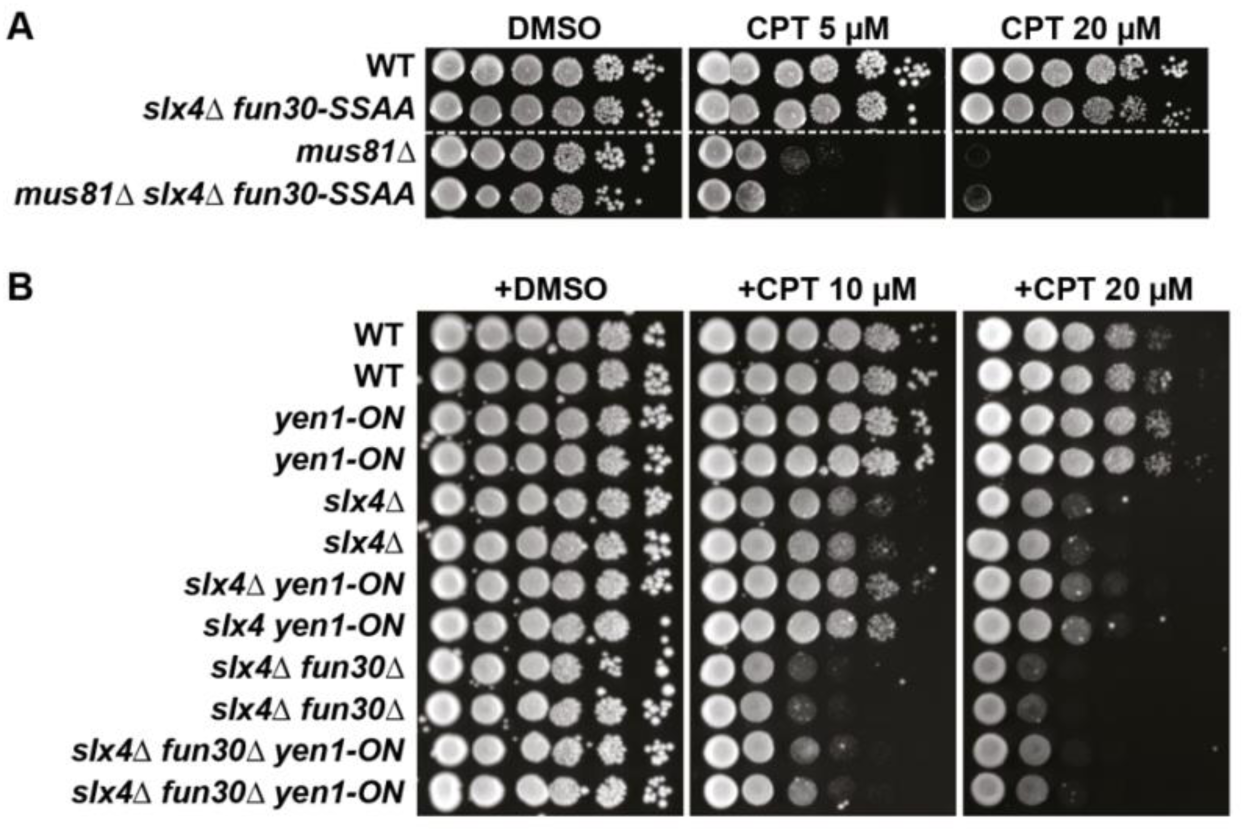
Yen1 uncontrolled activity does not compensate for the loss of checkpoint dampening. **A, B** – CPT sensitivity assay of indicated genotypes. 10-fold serial dilutions of yeast cells were spotted on YPAD growth plates containing increasing doses of CPT and incubated for 2 days at 30°C. Three biological replicates have been performed.

To test whether Mus81 can be activated in the absence of checkpoint dampening, we monitored the phosphorylation status of its regulatory subunit, Mms4, which is required to activate the endonuclease (64, 65). Mms4 phosphorylation, detected as an electrophoretic mobility shift by Western blot, normally occurs in the G2/M phase of the cell cycle and is dephosphorylated in the subsequent G1 (64, 65). As shown in **Figure 5B,C**, Mms4 phosphorylation appeared as soon as the cells reached G2/M (starting 90 minutes after release) for wild type and *slx4Δ* and *fun30Δ* single mutants. Phosphorylation levels decreased as cells entered a new cell cycle (at 180 minutes for wild-type cells and 240 minutes for single mutants). In contrast, *slx4Δ fun30Δ* double mutants progressed more slowly through S phase than wild-type cells and remained blocked in G2/M as described above (**Figure 5B**). Consequently, Mms4 hyperphosphorylation was delayed, starting 120 minutes after release and remained very high until the end of the time course experiment.

Together, these results indicate that Mus81 can be activated despite sustained checkpoint activation in the absence of both Slx4 and Fun30. This also indicates that the G2/M block imposed by checkpoint signaling is compatible with Mus81 activation. Thus, the CPT sensitivity of *slx4Δ fun30Δ* cells cannot be explained by a failure to activate Mus81 and is likely due to defects in other pathways of damage processing.

### Checkpoint hyperactivation negatively regulates DNA resection

The DNA damage checkpoint has been shown to inhibit DNA end resection, a prerequisite for DNA repair by HR (59, 60). In particular, Rad53 and Rad9 impede the two major pathways of long-range resection: Rad53 directly inhibits Exo1 via phosphorylation (66, 67) while Rad9 impedes the Sgs1-Dna2 pathway through a mechanism that remains unclear (Bonetti et al., 2015; Ferrari et al., 2015). We therefore hypothesized that dampening Rad53 activity via Rad9 displacement could be required for the initial resection of reversed forks and subsequent fork protection by HR in CPT-treated cells.

Consistent with this model, PFGE analysis revealed that loss of DNA resection in CPT-treated *exo1Δ sgs1Δ* cells led to extensive chromosome fragmentation (**Figure 6A**), as previously observed in cells lacking Rad52 (Pardo et al., 2020). Since Exo1 activity is directly inhibited by Rad53 phosphorylation, we next examined whether Exo1 was hyperphosphorylated in *slx4Δ fun30Δ* cells exposed to CPT. Exo1 electrophoretic mobility shifts were detected in both wild-type and *slx4Δ fun30Δ* cells exposed to CPT (**Figure 6B**). This shift was absent in cells expressing the *exo1-23A* allele, in which all known Rad53 phosphorylation sites were mutated to non-phosphorylatable residues (67), confirming that Exo1 mobility changes correspond to phosphorylation (**Figure 6B**). While Exo1 phosphorylation was modest and transient in wild-type cells, *slx4Δ fun30Δ* cells displayed a stronger and more persistent shift, suggesting that the loss of checkpoint dampening leads to sustained inhibition of Exo1 activity (**Figure 6B**).

**Figure 6.**
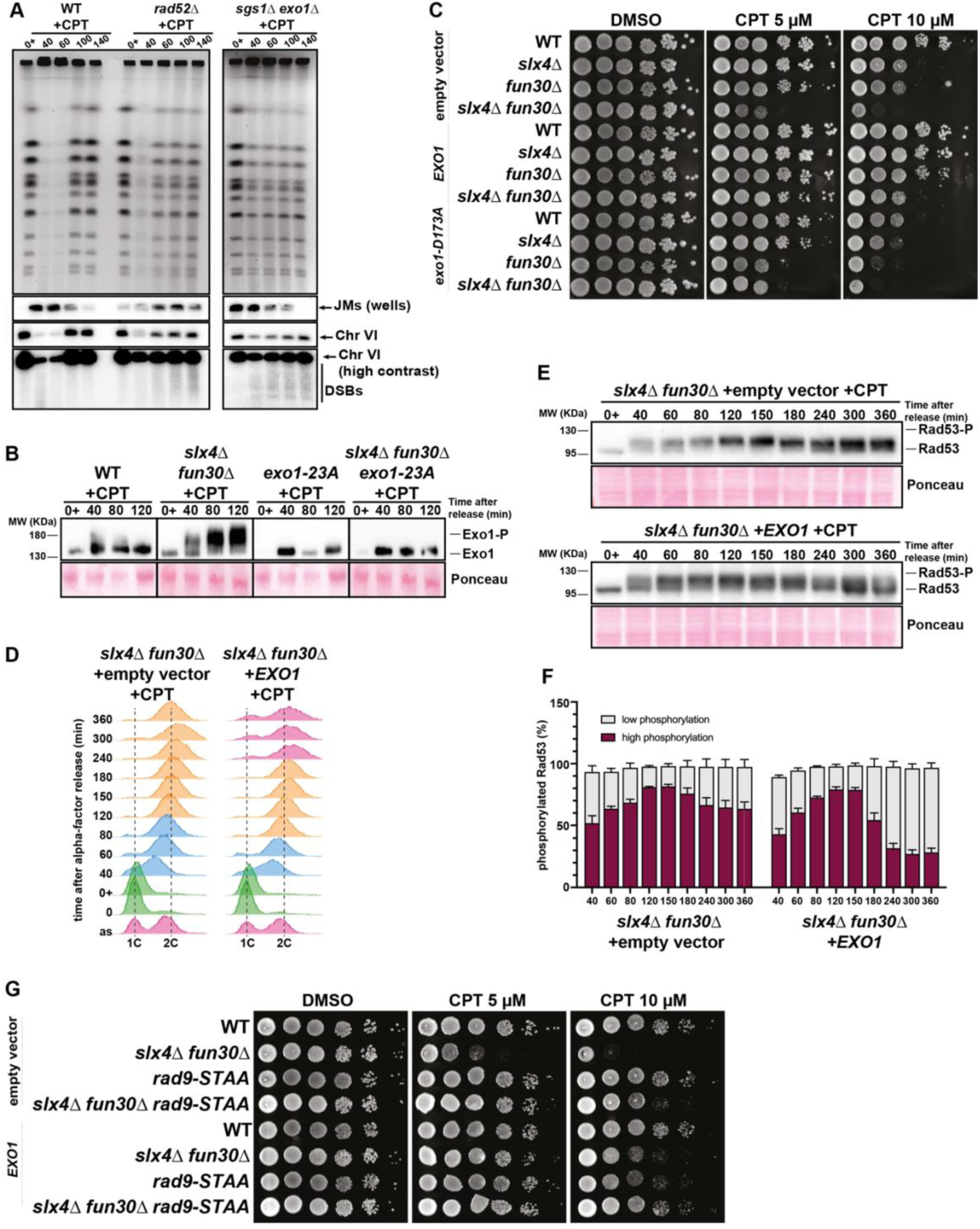
Checkpoint hyperactivation affects fork protection by inhibiting DNA resection. **A** – Analysis of chromosome breakage by pulsed-field gel electrophoresis (PFGE) in indicated genotypes. Cells were treated as in Figure 1B and collected at the indicated time points. DNA extracted in agarose plugs was analyzed by PFGE. Upper panel: agarose gel stained with ethidium bromide. Lower panels: Southern blot using a probe specific to chromosome VI. JMs, joint molecules accumulated in the gel wells. DSBs are revealed by the smeared signal detected below the chromosome VI band after overexposure of the hybridization signals. Two biological replicates have been performed. **B**– Exo1 phosphorylation analyzed by western blot using PhosTag reagent in indicated genotypes. Cells were blocked in G1 by using α-factor, treated with 100µM CPT for one hour (0+) in MPD +SDS medium and synchronously released into S-phase still in presence of drug. Cells were collected at the indicated time points and Exo1 was immunodetected with an anti-Myc antibody. Two biological replicates have been performed. **C, G** – CPT sensitivity assay of indicated genotypes. 10-fold serial dilutions of yeast cells were spotted on SC growth plates lacking tryptophane and containing increasing doses of CPT. Plates were incubated for 3 days at 30°C. Three biological replicates have been performed. **D** - Analysis of DNA content (1C, 2C) by flow cytometry of the experiments shown in panel E. **E**– Rad53 phosphorylation analyzed by western blot in wild-type and indicated mutants performed as in Figure 1B. Three biological replicates have been performed. **F** – Quantification of Rad53 phosphorylation showing the ratio of phosphorylated Rad53 (low and high)/total Rad53 observed by Western blot in indicated mutants. Data represent the mean ± SEM of three biological replicates.

*EXO1* overexpression has previously been shown to suppress CPT sensitivity and DSB resection defects caused by Fun30 deficiency (57). To determine whether resection defects could explain the CPT sensitivity of *slx4Δ* and *slx4Δ fun30Δ* mutants, we overexpressed various *EXO1* alleles in wild-type, *slx4Δ*, *fun30Δ*, and *slx4Δ fun30Δ* strains. Cells were transformed with either an empty vector, a vector carrying the wild-type *EXO1*, or the *exo1-D173A* nuclease-dead allele. As shown above, *slx4Δ* and *fun30Δ* mutants displayed additive sensitivity to CPT (**Figure 6C**). Overexpression of wild-type *EXO1*, but not the *exo1-D173A* mutant, fully suppressed the CPT sensitivity of both *slx4Δ* and *fun30Δ* single mutants and partially rescued the double mutant. Interestingly, overexpression of the *exo1-D173A* nuclease-dead mutant had a dominant-negative effect, exacerbating CPT sensitivity in wild type, *slx4Δ* and *fun30Δ* mutants. These results suggest that checkpoint hyperactivation, resulting from the loss of checkpoint dampening, negatively regulates DNA resection. Enhancing Exo1 resection activity is beneficial for cell survival and should therefore allow cell cycle resumption following CPT exposure.

We tested this hypothesis by monitoring cell cycle progression by flow cytometry and Rad53 phosphorylation by western blot in *slx4Δ fun30Δ* cells overexpressing *EXO1* (**Figure 6D,E**). In *slx4Δ fun30Δ* cells containing an empty vector, checkpoint hyperactivation persisted up to 360 minutes after release. In contrast, *EXO1* overexpression led to reduced Rad53 phosphorylation by 240 minutes, coinciding with partial cell cycle re-entry (**Figure 6E,F**). This phenotype is reminiscent of the *rad9-STAA* allele, which prevents checkpoint hyperactivation by disrupting the binding of Rad9 to Dpb11 (**Figure 3**). Indeed, *slx4Δ fun30Δ rad9-STAA* mutants showed a similar degree of CPT resistance to *slx4Δ fun30Δ* cells overexpressing *EXO1* (**Figure 6G**). Moreover, combining the overexpression of *EXO1* with *rad9-STAA* in *slx4Δ fun30Δ* cells produced a modest additive CPT resistance, suggesting that both recue strategies promote cell viability by different means.

Taken together, these results support a model in which downregulation of checkpoint activity restores fork resection and subsequent HR-mediated fork protection. Mec1 recruitment and activity may decrease at protected forks, decreasing in turn Rad53 and Chk1 phosphorylation and relieving the G2/M block.

### Characterization of CPT-dependent Top1ccs at a single locus

Our analyses thus far relied on methods that broadly indicate the presence of CPT-induced DNA damage throughout the yeast genome. To strengthen our conclusions, a more precise characterization of a specific damage site would be beneficial. However, genome-wide mapping of CPT-induced DNA damage sites in budding yeast genome has not yet been made. Of interest, Top1ccs has been shown to be specifically stabilized at the replication fork barrier (RFB) present in ribosomal DNA repeats (70), where it prevents conflicts between the DNA replication and transcription machineries. The Fob1 protein, which specifically binds to the RFB sequence, is responsible for the recruitment of Top1, which would be trapped at this site upon CPT treatment. Despite this, the endogenous rDNA RFB is not amenable to our study as it represents a natural barrier to DNA replication. In addition, the rDNA is largely refractory to checkpoint activation (71).

To overcome these limitations, we used a previously described system in which the rDNA RFB is integrated downstream of a replication origin (*ARS453*) in an orientation that allows fork passage, creating an ectopic RFB (eRFB) on chromosome IV (70). The RFB sequence was integrated alongside with a *KAN* resistance gene to select for positive integrations (**Figure 7E**; Bentsen et al., 2013). To increase the stability of Top1ccs, we used cells lacking Tdp1 and Wss1, two enzymes that act redundantly to remove Top1ccs in *S. cerevisiae* (5). Since *tdp1Δ wss1Δ* double mutant cells display growth defects even under unchallenged conditions (5), we constructed an auxin-inducible degron (AID) of Wss1 in *tdp1Δ* cells enabling conditional depletion of Wss1 in G1-arrested cells, before S-phase entry (**Figure 7A**).

**Figure 7.**
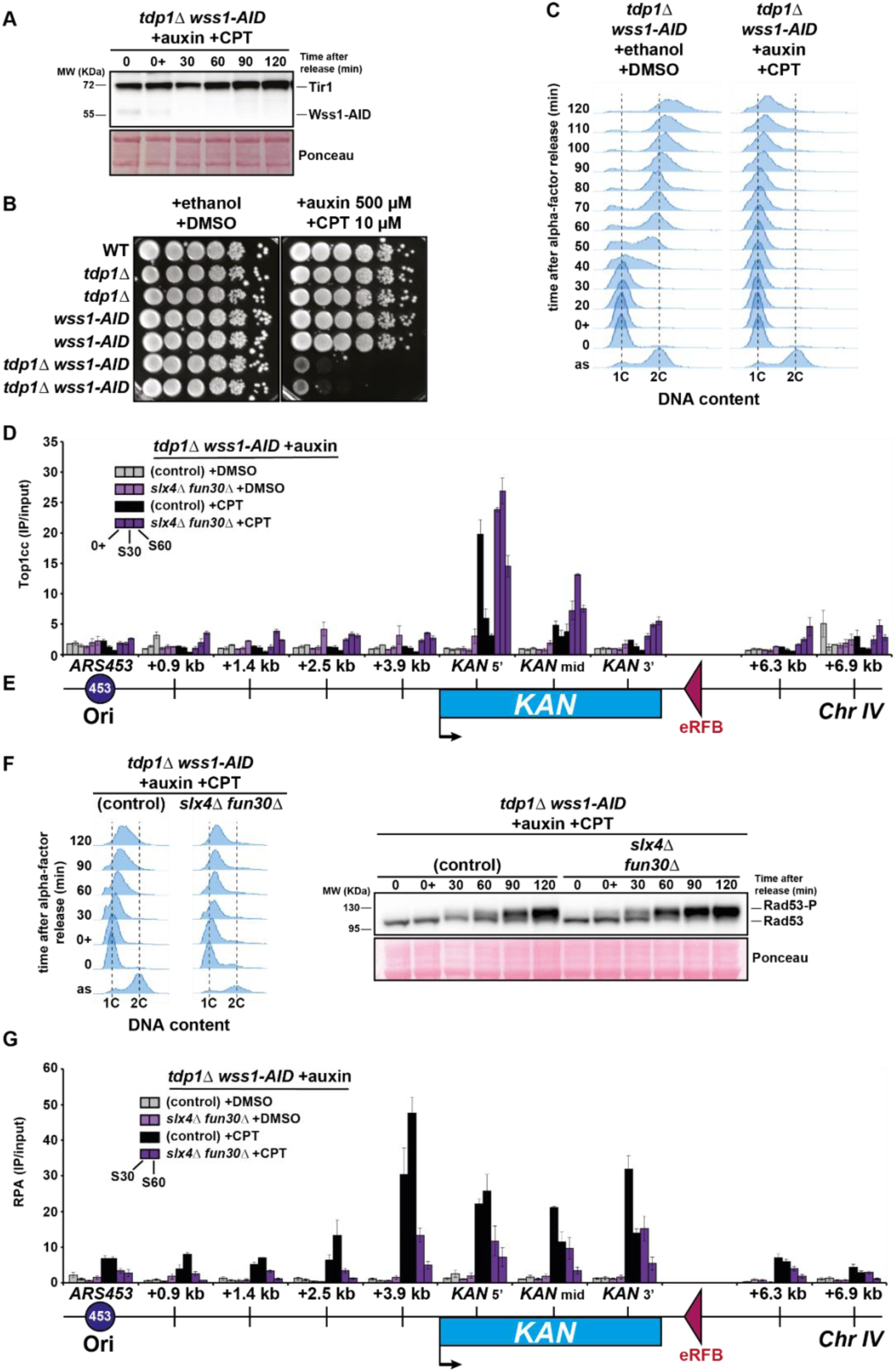
Loss of checkpoint dampening inhibits DNA resection at a specific Top1cc site in CPT-treated *tdp1Δ wss1-AID* cells. **A -** Wss1 degradation analyzed by western blot in *tdp1Δ wss1-AID* cells. Cells were blocked in G1 by using α-factor, treated with 1 mM auxin and 50µM CPT for one hour (0+) in MPD +SDS medium and synchronously released into S-phase still in presence of drugs. Cells were collected at the indicated time points and Wss1 and Tir1 were immunodetected with an anti-Myc antibody. Two biological replicates have been performed. **B** – CPT sensitivity assay of indicated genotypes. 10-fold serial dilutions of yeast cells were spotted on YPAD growth containing the indicated concentrations of auxin and CPT. Plates were incubated for 2 days at 30°C. Three biological replicates have been performed. **C** - Analysis of DNA content (1C, 2C) by flow cytometry in a time course experiment. Cells were collected at the indicated time points after being blocked in G1 phase with α-factor (0), treated with 1 mM auxin and 50 µM CPT or ethanol and DMSO for one hour (0+) in MPD +SDS medium and synchronously released into S phase still in presence of drugs. Three biological replicates have been performed. **D** –ChIP-qPCR analysis of Top1cc accumulation in the eRFB region in the indicated genotypes. Cells were blocked in G1 by using α-factor, treated with 1 mM auxin and 50µM CPT or DMSO for one hour (0+) in MPD +SDS medium and synchronously released into S-phase still in presence of drugs. Cells were collected at the indicated time points and processed for ChIP analysis without formaldehyde crosslinking. PCR primer pairs correspond to the regions upstream and downstream of the eRFB. Top1cc enrichment was normalized on a negative region for Top1 binding, not replicated in the experimental conditions. Mean and SEM correspond to two biological replicates. **E** – Schematic representation of the eRFB region on chromosome IV showing the relative positions of PCR primer pairs used in panel D. Primers are specific to *ARS453* and regions downstream of *ARS453* (indicated in kb) and to the *KAN* gene (3’, mid and 5’ regions). The scheme is not on genomic scale. **F**– Rad53 phosphorylation analyzed by western blot in control and *slx4Δ fun30Δ* mutant in *tdp1Δ wss1-AID* background. Cells were blocked in G1 by using α-factor, treated with 1 mM auxin and 50µM CPT for one hour (0+) in MPD +SDS medium and synchronously released into S-phase still in presence of drugs. Cells were collected at the indicated time points and Rad53 was immunodetected with an antibody against total Rad53. Flow cytometry profiles of DNA contents are also shown. Two biological replicates have been performed. **G** –ChIP-qPCR analysis of RPA enrichment in the eRFB region in the indicated genotypes. Cells were blocked in G1 by using α-factor, treated with 1 mM auxin and 50µM CPT or DMSO for one hour (0+) in MPD +SDS medium and synchronously released into S-phase still in presence of drugs. Cells were collected at the indicated time points and processed for ChIP analysis. PCR primer pairs correspond to the regions upstream and downstream of the eRFB. RPA enrichment was normalized on a negative region, not replicated in the experimental conditions. Mean and SEM correspond to two biological replicates.

**Supplementary Figure 7.**
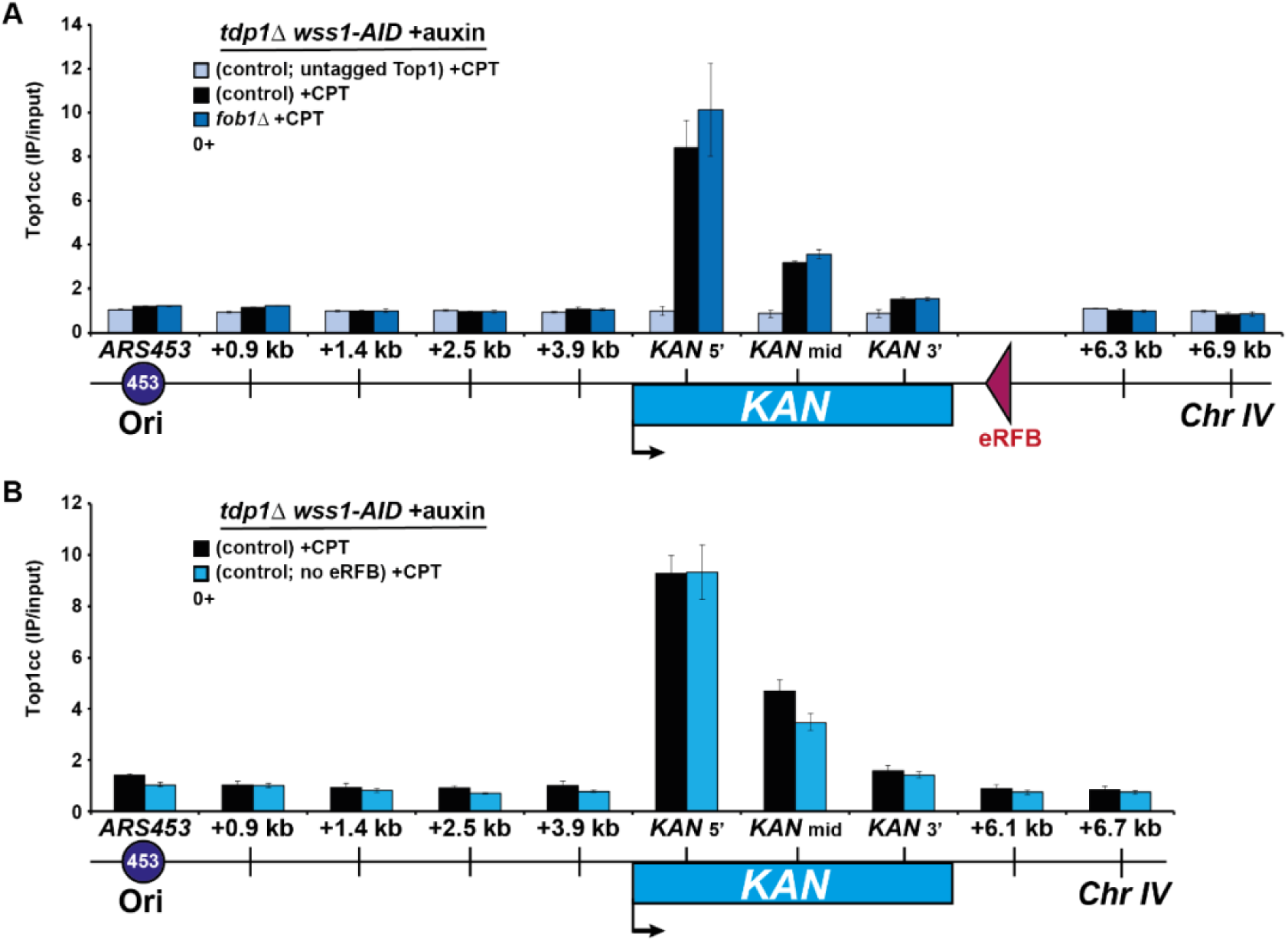
Top1ccs accumulation in the *KAN* gene in CPT-treated *tdp1Δ wss1-AID* cells does not depend on Fob1 nor the RFB sequence. **A, B -** ChIP-qPCR analysis of Top1cc accumulation in the *KAN* gene in *tdp1Δ wss1-AID* cells lacking FLAG tagged-Top1 (untagged Top1), the *FOB1* gene (*fob1Δ*) or the RFB sequence (no eRFB). Cells were blocked in G1 by using α-factor and treated with 1 mM auxin and 50µM CPT for one hour (0+) in MPD +SDS medium. Cells were processed for ChIP analysis. PCR primer pairs correspond to the regions upstream and downstream of the *KAN* gene. Top1cc enrichment was normalized on a negative region for Top1 binding. Mean and SEM correspond to three biological replicates.

In the presence of auxin, *tdp1Δ wss1-AID* cells showed increased CPT sensitivity, similar to that of *tdp1Δ wss1Δ* cells (**Figure 7B**; (5)). To further characterize this strain, we studied its cell cycle progression by flow cytometry. Cells were arrested in G1 with α-factor, incubated with auxin+CPT or ethanol+DMSO (control) for one hour prior to release into S phase. In contrast to control cells, those incubated with auxin+CPT progressed very slowly through S phase (**Figure 7C**). These results suggest that in the absence of Tdp1 and Wss1, the accumulation of Top1ccs in CPT-treated cells impedes DNA replication.

To determine whether Top1ccs accumulate at the eRFB, Top1 was tagged with 3 FLAG epitopes at its C-terminus and Top1-bound DNA was recovered by anti-FLAG immunoprecipitation. Importantly, no chemical crosslinking was used, as Top1 remains covalently linked to DNA after CPT treatment. We first examined Top1ccs distribution in G1-arrested *tdp1Δ wss1-AID* cells incubated for one hour either with auxin+DMSO or auxin+CPT. To quantify Top1-bound DNA, we designed several qPCR primer pairs spanning the eRFB locus, including *ARS453*, the *KAN* gene, and the region downstream of eRFB (**Figure 7E**). qPCR results showed that under auxin+DMSO conditions, only background levels of DNA were amplified, similar to those recovered from untagged Top1 immunoprecipitation (**Supplementary Figure 7A**). In contrast, a significant enrichment in Top1-bound DNA was observed upon CPT treatment in the *KAN* gene proximal to the eRFB, suggesting that Top1 is specifically trapped at this site in CPT-treated cells. This enrichment was unexpectedly stronger in the 5’ region of the *KAN* gene than in the 3’ region, despite the latter being closer to the RFB sequence.

Since Top1 recruitment to the endogenous rDNA RFB was shown to depend on Fob1, we repeated the experiment in *fob1Δ* cells (**Supplementary Figure 7A**). However, Top1 trapping at the *KAN* locus remained unchanged in the absence of Fob1. Moreover, similar results were obtained when the *KAN* gene was integrated at the same location without the eRFB (**Supplementary Figure 7B**). Altogether, these results indicate that although the eRFB fails to recruit Top1 at an ectopic location, the *KAN* gene inserted downstream of *ARS453* on chromosome IV represents a locus of choice to study CPT-induced DNA damage. Indeed, the strong expression of the *KAN* gene driven by the *TEF1* promoter may generate topological constraints that attract Top1.

Finally, we examined the fate of these Top1ccs following DNA replication. Cells arrested in G1 and treated with auxin and CPT were synchronously released into S phase, and Top1-bound DNA was monitored over time. We observed a progressive decrease in Top1cc levels within the *KAN* gene at 30 and 60 minutes after release (**Figure 7D**), suggesting a replication-dependent removal of Top1ccs that occurs independently of Tdp1 and Wss1.

### Loss of checkpoint dampening at a specific Top1cc site inhibits DNA resection during DNA replication

Then, we took advantage of this locus-specific assay to investigate how checkpoint dampening factors regulate locally the cellular responses to Top1ccs during S phase. We first examined Top1cc levels at the *KAN* gene in *tdp1Δ wss1-AID slx4Δ fun30Δ* cells treated with auxin and CPT. The levels of Top1ccs in G1 and after entry into S phase were similar to those detected in control cells (**Figure 7D**), indicating that Top1cc accumulation does not depend on Slx4 and Fun30. Next, we investigated the effect of Tdp1 and Wss1 loss on checkpoint activation in the absence of Slx4 and Fun30. To this end, G1-arrested cells were treated with auxin and CPT for one hour and were released into S phase in the presence of drugs. In control cells, Rad53 phosphorylation, monitored by western blot, was detected upon entry into S phase (t=30 min post release) and gradually increased at later time points (**Figure 7F**). Although the kinetics of Rad53 phosphorylation was similar in *slx4Δ fun30Δ* cells, the intensity of Rad53 phosphoshift was markedly higher, indicating stronger checkpoint activation. Given our prior results suggesting that Rad9 competes with Slx4 and Fun30 for Dpb11 binding, we hypothesized that this hyperactivation was due to at lesion sites in the absence of Slx4 and Fun30.

We next tested whether Rad9 stabilization inhibits DNA resection locally. To do so, we indirectly assessed DNA resection by monitoring the recruitment of the ssDNA-binding complex RPA by ChIP, as previously shown during hydroxyurea (HU)-induced replication stress (72). G1-blocked *tdp1Δ wss1-AID* cells were treated with auxin and CPT or DMSO as described above and RPA ChIP was performed 30 and 60 minutes after their release into S phase. In control cells, RPA was enriched at the *KAN* gene and upstream at *ARS453* +3.9 kb 30 minutes after release (**Figure 7G**). This enrichment increased at 60 minutes and extended to *ARS453* +2.5 kb. These results are consistent with the occurrence of DNA resection at the Top1cc sites and upstream region. Strikingly, RPA recruitment was markedly reduced in the in *slx4Δ fun30Δ* mutant (**Figure 7G**), indicating that DNA resection was inhibited in the absence of checkpoint dampening. Taken together, our results indicate that Rad9 displacement by Slx4 and Fun30 is important for allowing DNA resection at a specific Top1cc site.

## Discussion

In this study, we investigated the regulation of the checkpoint response to camptothecin (CPT), a Top1 poison that induce DNA damage and replication stress by trapping covalent Top1-DNA complexes (Top1ccs). Our data show that the cellular response to CPT-induced damage is strongly attenuated by Slx4-Rtt107 and Fun30, factors known to participate in DNA repair. This dampening mechanism operates through the displacement of Rad9, a key mediator of the DDC, from DNA damage sites by Slx4-Rtt107 and Fun30. Loss of checkpoint dampening leads to a strong inhibition of the cell cycle progression but is also associated with defective DNA end resection, which is required for protecting forks stalled by Top1-DNA crosslinks. These conclusions are further supported by our analysis of DNA replication through a specific locus containing Top1-DNA crosslinks induced by CPT.

As a classical model organism, the yeast *Saccharomyces cerevisiae* has been instrumental in advancing our understanding of eukaryotic cell cycle checkpoint pathways in response to DNA damage (DDR) (73, 74). However, since the discovery of CPT, our understanding of how yeast cells respond to this drug has remained very limited. Most analyses have focused on assessing the sensitivity of checkpoint mutants to chronic CPT exposure on solid media, largely because acute CPT treatment in liquid cultures produces minimal detectable effects on cell cycle progression (12, 35, 75). We previously demonstrated that this is mainly due to the low solubility of CPT in culture media and its inefficient uptake by yeast cells. By modifying the culture medium to increase this uptake, we revealed that CPT treatment of synchronized cultures induced delays in both S and G2/M phases (11). Here, we show that these cell cycle delays are abolished when the DNA damage response is inactivated. These results imply that the slowed S phase in CPT-treated cells is not primarily due to physical impediments to DNA replication but rather results from checkpoint signaling pathways.

This mechanistic insight explains why the S-phase delay is independent of the activation of the DRC branch of the S-phase checkpoint, which is mediated by Mrc1 and typically responds to replication impediments such as dNTP depletion by HU or DNA alkylation by MMS (27). Instead, the response is mediated by the DDC, consistent with a response triggered directly at DNA lesion sites. We have previously shown that Rad9-dependent signaling modulates DNA replication dynamics by restricting both origin firing and fork progression (18), a regulatory mechanism also observed in human cells treated with CPT (76). The yeast model remains therefore a powerful tool to further dissect the molecular basis of this replication control mechanism.

We further demonstrate in this study that the checkpoint response to CPT-induced DNA damage is strongly attenuated by the coordinated actions of Slx4-Rtt107 and Fun30. Checkpoint dampening by Slx4-Rtt107 has been well documented in the context of replication stress induced by HU and MMS (31). Slx4 and Rtt107 compete with Rad9 for binding to Dbp11 and γH2A(X), thereby displacing Rad9 from damage sites and reducing activation of the checkpoint effector kinase Rad53. Our findings reveal an additional layer of regulation through the involvement of Fun30. Specifically, we show that loss of the ability of Fun30 to bind Dbp11 further increase the checkpoint response in the absence of Slx4. Since we also observed that the chromatin remodeling activity of Fun30 is not required for checkpoint dampening, we propose that Fun30 displaces Rad9 only by competing for the binding to the same domain of Dpb11 (52) rather than through the eviction of modified nucleosomes, to which Rad9 can bind (77). In the fission yeast *Schizosaccharomyces pombe*, the Fun30 homolog Fft3 is required for the restart of collapsed replication forks independently of its ATPase activity (78). Although the lack of checkpoint activity does not affect fork restart at this site-specific replication fork barrier (79), the impact of checkpoint hyperactivation has not been investigated. It is therefore plausible that Fft3 promotes fork restart by dampening checkpoint activity, in a manner analogous to the function we describe for Fun30 in budding yeast. Local checkpoint attenuation can also be mediated by the PP2A phosphatase, which impedes the interaction between Dpb11 and the 9-1-1 complex (80). Moreover, while this manuscript was in preparation, the Zhao lab reported that Rad9 can also be displaced from DNA damage sites by the Mms22-Rtt107 complex by promoting the proteasomal degradation of Rad9 (81). This pathway acts in parallel with the Slx4-Rtt107 dampening pathway, underscoring the importance of multiple, partially redundant mechanisms for downregulating the DNA damage checkpoint.

We noticed that checkpoint dampening in response to CPT resulted in markedly reduced Rad53 activation in wild-type cells compared to treatment with HU or MMS. A simple explanation for the low-level activation is that CPT induces relatively few DNA lesions, likely due to its poor uptake by yeast cells, even at high concentrations. However, our findings suggest an alternative possibility in which checkpoint dampening is particularly important for the maintenance of CPT-arrested forks. Indeed, we show that in the absence of Slx4 and Fun30, the DDR becomes hyperactivated, despite the presence of several phosphatases acting on Rad53 (82, 83). This hyperactivity correlates with impaired HR in response to CPT. We have previously proposed that HR promotes DNA replication under CPT-mediated stress by protecting stalled forks (11). Our current data indicate that excessive checkpoint activation interferes with this process by impairing fork resection, a prerequisite for HR. Notably, we found that in *slx4Δ fun30Δ* cells exposed to CPT, the number of cells bearing Rfa1 foci persisted over time. Since Rfa1 is part of the RPA complex that binds single-stranded DNA (ssDNA), these data could be interpreted at first glance as evidence of excessive ssDNA generation due to over-resection. However, similar phenotypes have been observed in cells defective in processive resection (84), suggesting an alternative explanation. Supporting this, we observed increased phosphorylation of the Exo1 nuclease in CPT-treated *slx4Δ fun30Δ* cells, a modification that has been described to inhibit its resection activity (66, 67). Furthermore, our analysis of a CPT-induced Top1-DNA crosslink at a defined genomic site revealed impaired RPA recruitment during replication in the absence of Slx4 and Fun30. In contrast, RPA accumulated at this locus in control cells, presumably due to the timely resection of nascent DNA at reversed forks. In *slx4Δ fun30Δ* cells, this resection defect is associated with a stabilization of Rad9 at the locus. Rad9 has been shown to inhibit DNA resection at DSBs (85) and during post-replicative repair of MMS-induced DNA lesions (30). Our study extends this role to replication forks encountering Top1ccs. Importantly, long-range resection by Exo1 is not required for the restart of damaged forks in *S. cerevisiae* (86) or in *S. pombe* (79, 87), raising questions about the necessity for extensive resection at forks encountering Top1ccs. Moreover, replication fork restart does not appear to occur in cells exposed to CPT (11). We therefore propose that reversed forks formed at damage sites require protection from nucleolytic degradation until they can be rescued by converging forks. Paradoxically this protection may depend on the promotion of Exo1 activity, which is impaired when checkpoint signaling is not properly attenuated.

We have previously shown that the Mus81 endonuclease is required during G2/M to promote replication completion in CPT-treated cells (11). Other studies have suggested that the S-phase checkpoint could restrict Mus81 activity during HR-mediated repair, as premature Mus81 engagement has been observed in checkpoint-deficient mutants (62). It had also been proposed that checkpoint hyperactivation in the absence of Slx4 and the Pph3 phosphatase, which targets activated Rad53, prevents the resolution of HR intermediates by Mus81 (61). In this study, our genetic analyses show that loss of checkpoint dampening by Slx4 and Fun30 is not equivalent to loss of Mus81 in response to CPT-induced DNA damage, indicating that the function of Slx4 and Fun30 is distinct from enabling Mus81 engagement. Supporting this, we found that Mus81 activation by Mms4 phosphorylation occurs even when the checkpoint is hyperactivated in *slx4Δ fun30Δ* cells blocked at the G2/M transition. We conclude that the DNA damage response is unlikely to directly interfere with Mus81 processing of HR intermediates. However, the absence of signaling in checkpoint mutants would allow premature entry into G2/M and Mus81 activation, leading to untimely processing of DNA structures, increasing the risk of chromosomal rearrangements.

Dampening of the DNA damage response in mammalian cells by factors related to yeast Slx4-Rtt107 and Fun30 has not yet been described. While SLX4, PTIP, and SMARCAD1 have been identified as homologs of Slx4, Rtt107, and Fun30, respectively, their role in modulating the checkpoint response has not been investigated. Nonetheless, it is well established that 53BP1, the mammalian ortholog of yeast Rad9, inhibits DSB end resection and dictates the choice of repair pathways (88). Like Rad9, 53BP1 binds to γH2AX (89) and acts as a mediator of ATR-CHK1 pathway in response to DNA damage (90, 91). Both functions require the interaction of 53BP1 with TOPBP1, the mammalian ortholog of yeast Dpb11 (30, 90, 91). The BRCA1-BARD1 ubiquitin ligase complex counteracts the inhibition of resection by 53BP1 (88), not only by directly stimulating the activity of nucleases (92), but also by displacing 53BP1 from break sites. This latter function depends on the chromatin remodeling activity of SMARCAD1 (57, 93), which also interacts with TOPBP1 (52). TOPBP1 thus emerges as a central regulator, linking ATR activation during S phase to the control of DNA end resection (23, 30, 94). Under replication stress, SMARCAD1 also interacts with PCNA at replication forks and facilitates the eviction of 53BP1-bound nucleosomes (95). This leads to increased recruitment of BRCA1-BARD1 to TOPBP1, promoting fork protection and restart (30, 95). Collectively, these data suggest that 53BP1 and SMARCAD1, like their yeast counterparts Rad9 and Fun30, exert opposing roles in DNA end resection and influence replication fork protection in mammalian cells. Whether the ability of 53BP1 to inhibit resection is mechanically linked to checkpoint signaling, and whether SMARCAD1 suppresses checkpoint activation by displacing 53BP1 remains to be explored.

Altogether, our results support the mechanistic model presented in **Figure 8**. We posit that replication forks undergo reversal upon encountering Top1ccs, a phenomenon previously observed in CPT-treated yeast cells by electron microscopy (10, 12). Fork reversal could be driven by the accumulation of topological constraints resulting from Top1 inhibition (96) and may provide more space for Top1cc removal. In the absence of Tdp1 and Wss1, Top1cc removal would require Top1 debulking by the proteasome (97) and removal of the phosphotyrosine bond by an endonuclease. Several 3’ flap endonucleases such as Mus81, Rad1^XPF^ and Mre11 have been implicated in this activity in the absence of Tdp1 (98–102). Endonucleolytic cleavage upstream of the phosphotyrosine bond would generate a ssDNA gap, which would be rapidly coated with RPA. This in turn would recruit the sensor checkpoint kinase Mec1, which phosphorylates nearby H2A histones, and promotes loading of the 9-1-1 complex and its Dpb11 interactor at ss/ds 5’ DNA junctions. Binding of Rad9 to both Dpb11 and γH2A(X) would stabilize it at the lesion site and promote Rad53 activation. Rad53 activation would then impede DNA resection by phosphorylating the Exo1 nuclease (66, 67). Checkpoint dampening by Slx4-Rtt107 and Fun30 displaces Rad9 and relieves Exo1 inhibition, enabling resection to proceed. The fact that RPA binding was detected at the Top1cc site and the upstream region indicates that ssDNA was generated upstream of the lesion site and/or at the reversed fork, which share the same DNA sequence. Fork resection would promote the HR-mediated invasion of the parental strand by the regressed arm. We propose that this structure provides protection against further degradation of nascent strands, a mechanism reminiscent of the protection of chromosome ends by telomere looping (103), until this HR intermediate merges with a converging fork (**Figure 8**) (11). The site-specific CPT-induced Top1cc system proved invaluable in constructing this model. Further use of this system will undoubtedly deepen our understanding of checkpoint dampening at the molecular level and, more generally, of the cellular mechanisms that maintain genome integrity in response to Top1ccs and other DNA-protein crosslinks.

**Figure 8.**
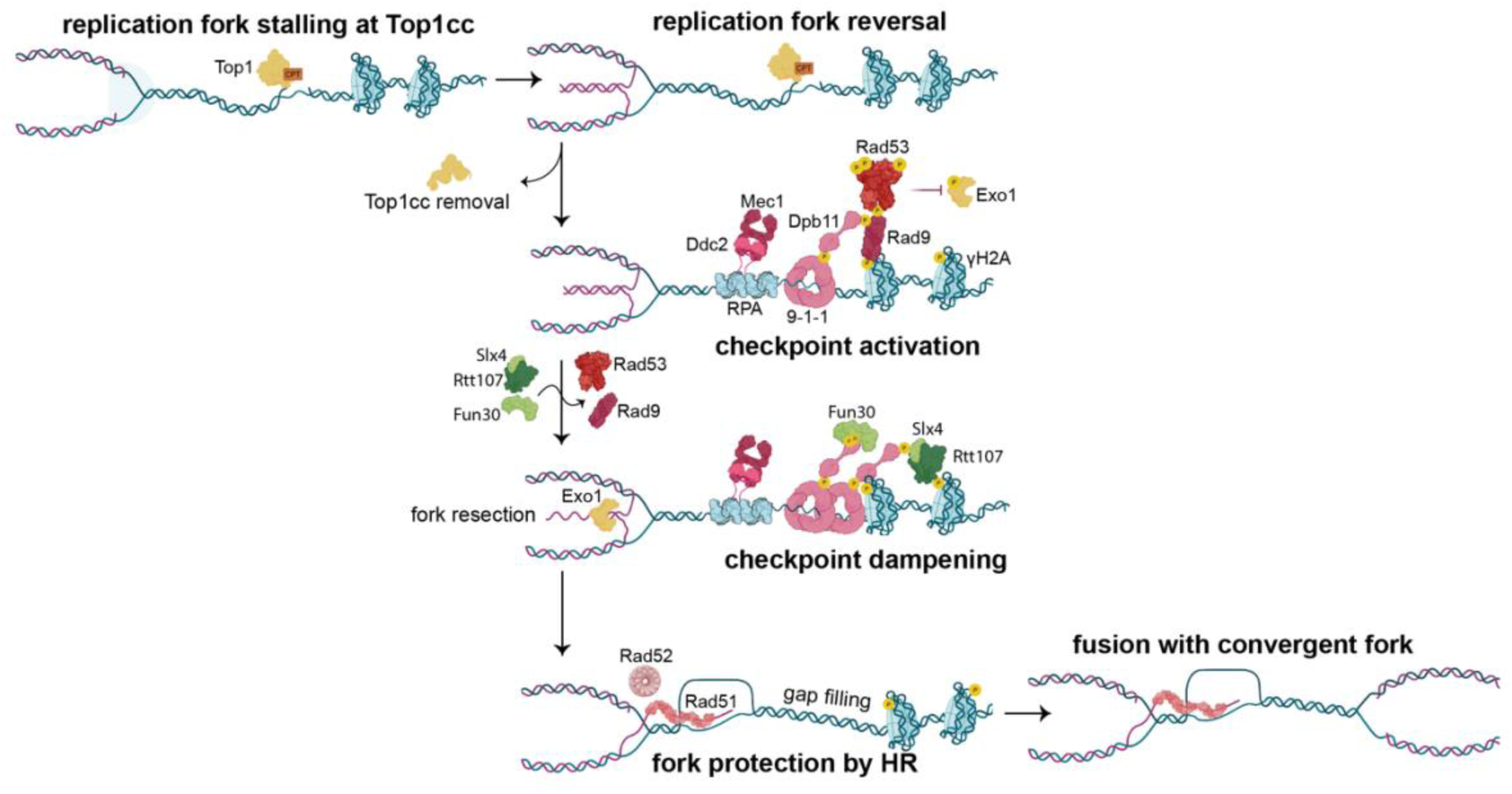
Proposed model for the function of checkpoint dampening in response to Top1-DNA crosslinks during DNA replication. See text for details.

## Data availability

The data underlying this article are available in the article and in its online supplementary material. Raw data will be shared upon reasonable request to the corresponding authors.

## Supplementary Data

Supplementary Data are available at *NAR* Online.

## Acknowledgements

We thank Boris Pfander, Bertrand Llorente, Miguel Blanco, Lotte Bjergbaek, Monica Segurado and Lorraine Symington for strains and plasmids. We also thank Corrado Santocanale for the Rad53 antibody. We thank the members of the Pasero laboratory for discussions and Yea-Lih Lin for critical comments on the manuscript. We acknowledge the imaging facility MRI, a member of the national infrastructure France-BioImaging, supported by the French National Research Agency [ANR-10-INBS-04, Investissements d’avenir].

## Author Contributions

Conceptualization, MC, TB, PP and BP; Methodology, MC, TB, AB, PP and BP; Supervision: PP and BP; Project Administration, PP and BP; Funding Acquisition, MC, PP and BP; Writing – Original Draft, MC and BP; Writing – Review & Editing, MC, PP and BP.

## Fundings

This work was supported by funds from the Ligue Contre le Cancer (équipe labélisée) and the Agence Nationale de la Recherche [ReSPoND] to PP and from the Fondation ARC pour la Recherche sur le Cancer [ARCPJA22020060002119] to BP. MC doctoral funding was supported by the Université de Montpellier and the Ligue Contre le Cancer [IP/SC-18293].

## Declaration of Interests

The authors declare no competing interests.

## Notes

### Competing Interest Statement

The authors have declared no competing interest.

